# Time-dependent computational model of post-traumatic osteoarthritis to estimate how mechanoinflammatory mechanisms impact cartilage aggrecan content

**DOI:** 10.1101/2024.10.08.617186

**Authors:** ASA Eskelinen, JP Kosonen, M Hamada, A Esrafilian, C Florea, AJ Grodzinsky, P Tanska, RK Korhonen

## Abstract

Degenerative musculoskeletal diseases like osteoarthritis can be initiated by joint injury. Injurious overloading-induced mechanical straining of articular cartilage and subsequent biological responses may trigger cartilage degradation. One early sign of degradation is loss of aggrecan content which is potentially accelerated near chondral lesions under physiological loading. Yet, the mechanoinflammatory mechanisms explaining time-dependent degradation in regions with disparate mechanical loading are unclear and challenging to assess with experiments alone. Here, we developed computational models unraveling potential mechanisms behind aggrecan content adaptation. Incorporating mechanical strain-driven cell damage and downstream proteolytic enzyme release, fluid flow-driven aggrecan depletion, and fluid pressure-stimulated regulation of aggrecan biosynthesis, the models agreed with experiments and exhibited 14%-points greater near-lesion aggrecan loss after 12 days of physiological loading compared to without loading. This significant advancement in mechanistic understanding incorporated into cartilage adaptation model can help in development and guidance of personalized therapies, such as rehabilitation protocols and tissue- engineered constructs.

## 1 Introduction

Post-traumatic osteoarthritis (PTOA) is a degenerative musculoskeletal disorder that develops over time following an injury to the knee joint. Injurious overloading and damage to articular cartilage are common in sports, rendering PTOA the predominant osteoarthritis phenotype in young active population^1,2^. In addition to visible tissue damage, mechanical insults and excessive mechanical loading may activate cell receptors and signaling pathways or even harm the cartilage cells which respond to this insult with catabolic enzyme production^3–5^. Defined as mechanoinflammation^6^, this cascade leads to compositional changes such as loss of cartilage aggrecan content^7,8^, an early sign of PTOA. However, it is yet not understood in which tissue regions and to what extent the underlying mechanoinflammatory mechanisms affect aggrecan content adaptation over time. Since evaluating the impact of different pathomechanisms experimentally is challenging, efforts could be reinforced by computational modeling to identify effective therapeutic solutions in halting PTOA progression.

The disruption of tissue homeostasis in early-stage PTOA has been studied experimentally using *ex vivo* cartilage explant cultures^5,7–14^. Following a controlled injurious overloading in (un)confined compression^15,16^ or drop-tower setups^11,17^, tissue-level catabolic responses such as substantial decreases in cell viability^5,7,18^, extracellular matrix (ECM) proteoglycan content (PG, such as aggrecan)^8,9,14,19^, and biosynthesis of new aggrecan^10,14,20^ have been reported within the first 1–3 days. The injurious compression can excessively deform and shear the ECM^18^ and increase the hydrostatic pressure rapidly to hyper-physiological levels^21–23^. Depending on the cellular microenvironment and the level of stress or strain (rate)^24–28^, cells may stay viable, undergo phenotypical changes (such as hypertrophy^29^), respond by exhibiting different types of cell damage (mitochondrial dysfunction and oxidative stress^11,13,29^), or die (apoptosis, necrosis)^18,28,30–32^. Cell damage alters cellular function that entails upregulation of proteolytic activity, *e.g.*, release of aggrecan-degrading aggrecanases. Aggrecanase activity has been detected *ex vivo*^4,14,33,34^ and aggrecanase-related loss of aggrecan has been observed *in vivo*^35^ within 1–2 days following injury.

Cyclic loading at normal physiological levels is beneficial for overall cartilage health. It promotes moderately varying fluid pressure over time, interstitial fluid flow, and solute transport^36,37^, resulting in anabolic responses and accelerated aggrecan biosynthesis compared to static loading in explant cultures^38–42^ and *in vivo*^43^. Yet, cyclic loading can also have locally degradative effects near lesions, where tissue (shear) stress/strain can rise to excessive levels, resulting in cell damage and subsequent aggrecan loss^7,44–47^. Physiological loading may also promote the transport of aggrecan fragments through damaged lesion surfaces via fluid flow^7,48^.

Computational cartilage adaptation models have been developed to gain an understanding of the link between difficult-to-measure intra-tissue mechanical responses and experimental tissue alterations. Previous mechanistic modeling frameworks with either injurious^3,49^ or cyclic loading^7,50^ have connected loss of matrix proteins (PGs, collagen) to shear or compressive strain (rate) and tensile stress^7,26,27,51^. These models also implemented the decrease of cartilage load-bearing capability by updating permeability, matrix stiffness, or swelling/osmotic properties^27,51–53^. Models incorporating cell-level responses^49,54,55^, chemo–mechano–biological insights^3^, and possible pro-anabolic effects^3,53^ are presently few but important advancements in numerical estimation of mechanoinflammation. However, understanding how tissue-level mechanical loading alters cell viability/function and how that reflects on experimentally observed, spatio-temporal adaptation of aggrecan content is lacking.

Our aim was to develop a mechanistic modeling framework to explain localized and time-dependent (12-day) aggrecan loss in young bovine knee cartilage predisposed to injurious loading (INJ), physiological cyclic loading (CL), and their combination (INJ+CL). The major novel aspect the framework is equipped with is the estimates of cartilage cell responses to PTOA-triggering mechanical loading, and how the ensuing cell-driven mechanoinflammation regulates spatial aggrecan content in the early disease stages. Intrigued by previous explant culture studies reporting depth-wise aggrecan loss and its worsening near chondral lesions with INJ+CL^7,44^ (Fig. 1AB), we hypothesized that CL would induce heterogeneous strain and fluid flow fields with functional importance for cartilage cell and aggrecan distributions^22,32^. To replicate experimental aggrecan loss, we built control, INJ, CL, and INJ+CL models in a step-by-step fashion, adding mechanoinflammatory cartilage adaptation mechanisms as necessary (cell damage with aggrecanase release, fluid flow-driven depletion of aggrecan, fluid pressure-stimulated acceleration of aggrecan biosynthesis; Fig. 1C), and performing sensitivity analyses.

**Fig. 1.**
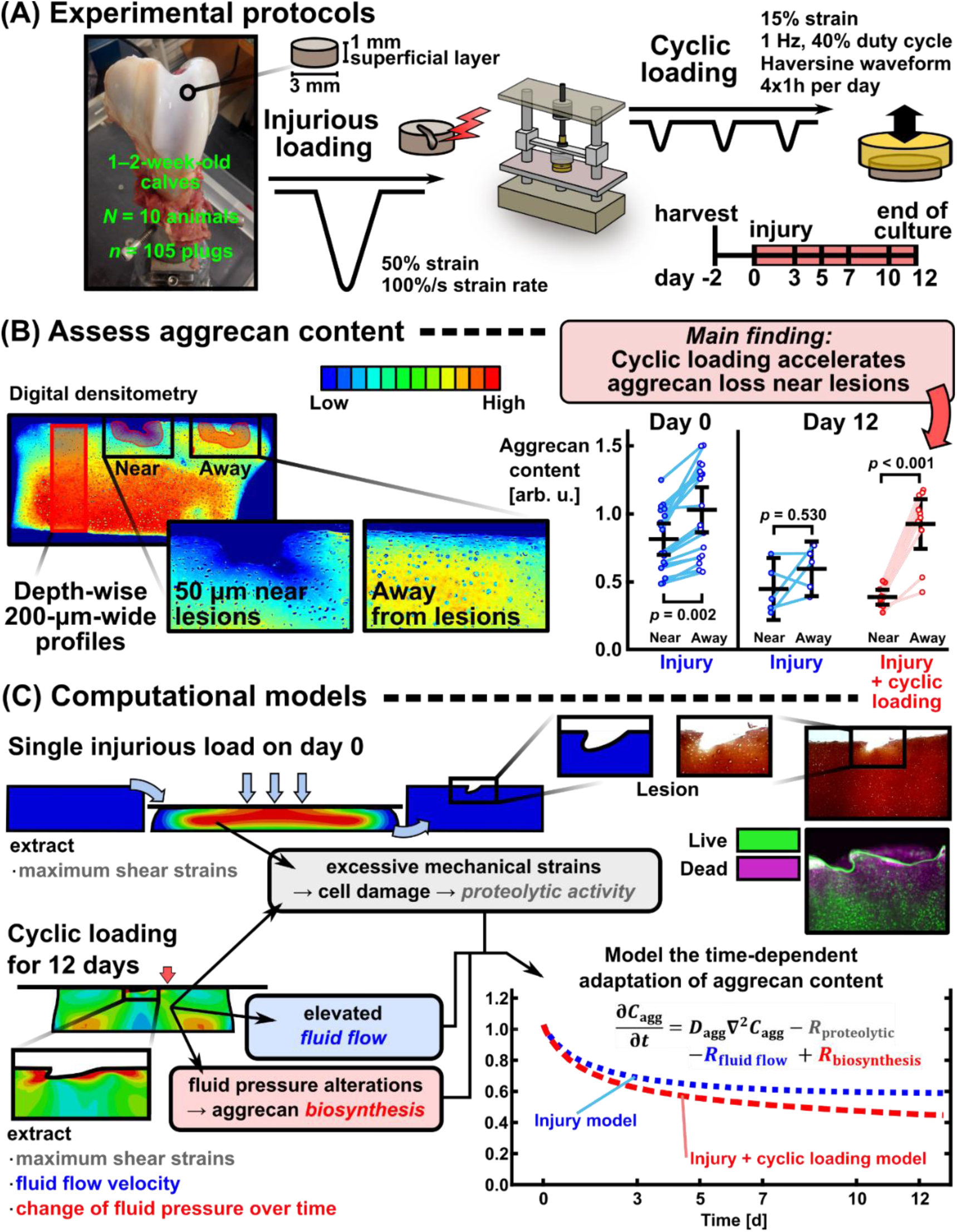
Study workflow. **(A)** In the experiments, cartilage explants were subjected to single injurious load–unload cycle followed by up to 12 days of physiological cyclic loading. **(B)** Cartilage aggrecan content was assessed from Safranin-O-stained sections with digital densitometry in depth-wise profiles and in regions near and away from chondral lesions. The plots with near and away from lesion data represent mean ± 95% confidence intervals, and linear mixed effects model with Bonferroni adjustment. The day 0 values are pooled from two previous datasets^7,44^. **(C)** Computational finite element models included single injurious loading followed by physiological cyclic loading. After implementing three time-dependent, mechanobiological cartilage adaptation mechanisms (proteolytic activity upregulated by damaged cells in highly sheared regions, fluid flow-driven transport of aggrecan fragments out of tissue, and accelerated aggrecan biosynthesis due to fluid pressure changes over time), the injury and cyclic loading model replicated accelerated aggrecan loss near lesions compared to away from lesions, as observed in the experiments. Lesion formation was not modeled in this study

## 2 Results

### 2.1 Cyclic loading accelerates experimental aggrecan loss near lesions

In the experiments, the aggrecan content within 50 µm from lesion edges was significantly lower than away from lesions on the day of injury (*p*=0.002, linear mixed effects (LME) model with Bonferroni adjustment, Fig. 1B). On day 12, the aggrecan contents near and away from lesions in the INJ group were similar (*p*=0.530). In the INJ+CL group, the near-lesion aggrecan content was significantly lower than away from lesions on day 12 (*p*<0.001, Fig. 1B).

### 2.2 Computational models of mechanoinflammation reproduced the injury-related through-depth aggrecan loss as well as the locally detrimental and beneficial effects of physiological loading

After implementing the initial day-0 aggrecan content (Fig. 2A), the computational model with intact geometry was first calibrated using day-12 depth-wise aggrecan content data from free-swelling control (CTRL) conditions. This model included free diffusion of aggrecan out of the tissue, resulting in an ∼8–25% decrease in aggrecan content in contrast to ∼10–30% mean decrease in the experiment at normalized depths of 0–40% (0%=surface, 100%=bottom of the cartilage explant) on day 12 compared to day 0 (Fig. 2B). The INJ model included a single injurious load–unload cycle; in regions of excessive shear strain at the time of maximum compression, ‘healthy’ cells were assumed to switch their phenotype to ‘damaged’ cell (only these two cell populations were implemented). Subsequently, the damaged cells released aggrecanases, promoting enzymatic activity for 12 days. This approach captured the substantial decrease in depth-wise aggrecan content in INJ compared to CTRL^7,44^ (Fig. 2BC). The CL models with physiological loading for 12 days included the following mechanisms: shear strain-driven cell damage (and, thus, localized enzymatic activity), depletion of aggrecan fragments out of the tissue in regions of high fluid flow velocity, and accelerated of biosynthesis of new aggrecan molecules in regions of moderate fluid pressure changes over time. The resulting depth- wise aggrecan contents in intact regions especially at normalized depths of 0–40% were higher in both the CL and INJ+CL models compared to the INJ, as observed in the experiments (Fig. 2DE).

**Fig 2.**
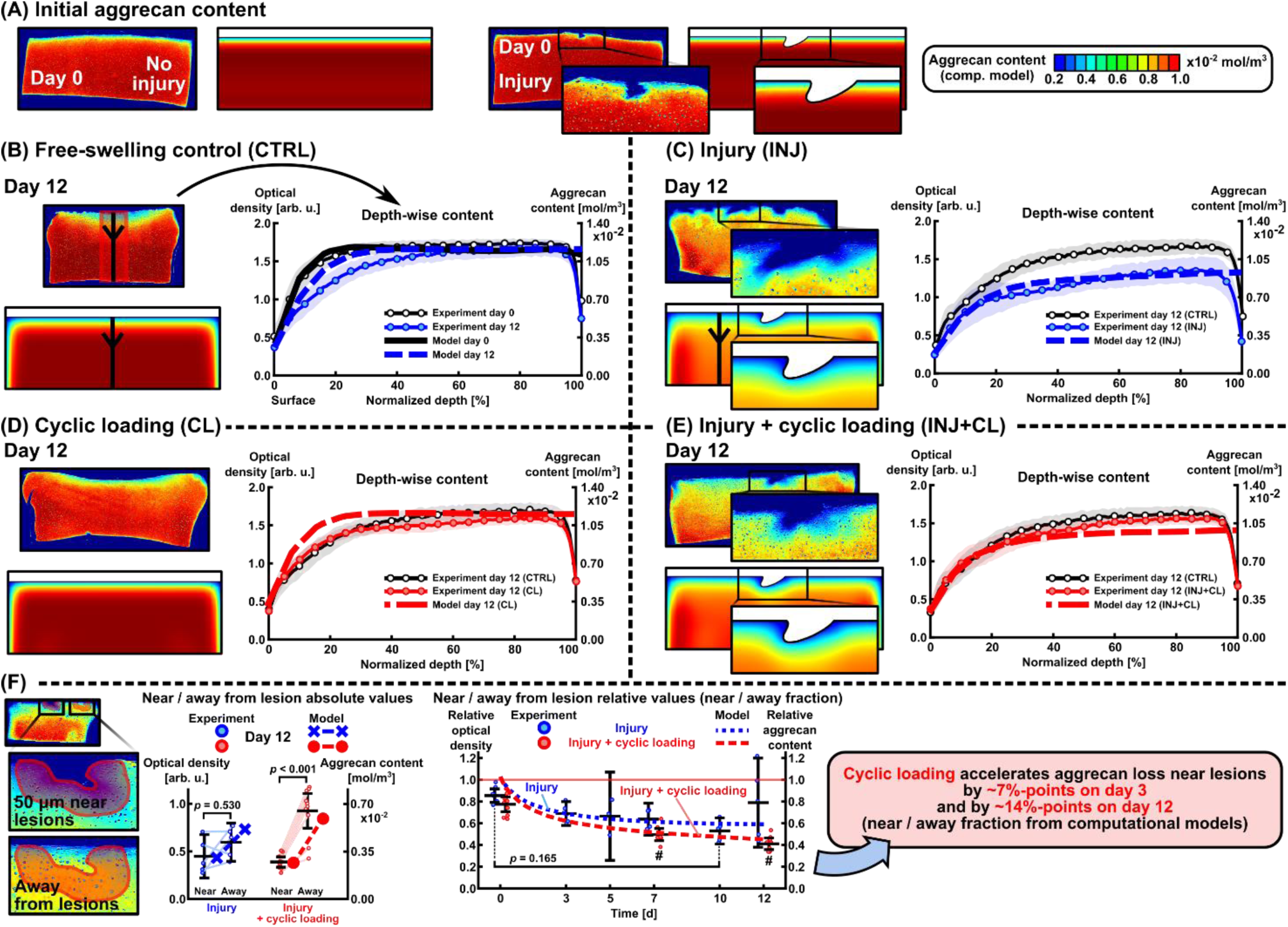
The calibrated computational model reproduced the experimental findings of aggrecan content adaptation depth-wise, and near and away from chondral lesions over 0–12 days. **(A)** Initial aggrecan distribution at the beginning of the culture and simulation. Depth-wise aggrecan content away from lesions in **(B)** free-swelling control, **(C)** injurious loading, **(D)** cyclic loading, and **(E)**injurious and cyclic loading experimental groups and computational models. **(F)** Numerical estimates of near and away from lesion aggrecan contents agreed with experiments in absolute (left panel) and relative terms (right panel; near/away fraction; value <1 means more aggrecan lost near lesions compared to away). The plots represent mean ± 95% confidence intervals, and linear mixed effects model with Bonferroni adjustment. **#** marks *p* < 0.001 compared to day 0 (right panel)

The near-lesion aggrecan content was lower than the away-from-lesion content in both the INJ and INJ+CL models on day 12 (Fig. 2F, left panel). The difference between near and away aggrecan contents was greater in the INJ+CL model (0.21 ⋅ 10^−2^ mol ⋅ m^−3^) than in the INJ model (0.32 ⋅ 10^−2^ mol ⋅ m^−3^) on day 12, consistent with the experiments (Fig. 2F, left panel). Over time, the relative near/away from lesion aggrecan content decreased more rapidly in the INJ+CL compared to the INJ model. For the INJ model, the relative aggrecan content was 0.689 on day 3 (experiment: 0.690±0.110 (95% confidence interval)), 0.614 on day 7 (0.638±0.147), 0.596 on day 10 (0.530±0.120, *p*=0.165 compared to day 0 (0.855±0.062)), and 0.591 on day 12 (0.790±0.410, Fig. 2F, right panel). For the INJ+CL model, the relative aggrecan content was decreased from 0.622 on day 3 to 0.516 on day 7 (experiment: 0.496±0.056, *p*<0.001 compared to day 0 (0.775±0.068)) and to 0.452 on day 12 (experiment: 0.411±0.053, *p*<0.001 compared to day 0, Fig. 2F, right panel). Cyclic loading accelerated aggrecan loss by 6.7%-points on day 3 (near *vs*. away fraction INJ: 0.689, INJ+CL: 0.622) and by 13.9%-points on day 12 (INJ: 0.591, INJ+CL: 0.452, Fig. 2F, right panel).

### 2.3 Injurious loading may increase catabolic cell activity via excessive mechanical shear strains and rapid fluid pressure changes

The modeled injurious loading triggered whole-thickness loss of healthy cells^7,14,15,44,56,57^ on day 0, observed as ∼30–45% increase in non-healthy (damaged) cells^5,54,56^ in the computational INJ model (Fig. 3A, right panel). Various literature-supported biomechanical mechanisms for triggering the injury-associated cell damage and subsequent catabolic cell activity (aggrecanase release) were investigated^3,22,46,56–59^: maximum shear strain (damage initiation threshold: 40%^48,56,60^, Fig. 3AB), rapid change of fluid pressure over the time of injurious compression (threshold: 80 MPa ⋅ s^−1^ ^22,59,61^, the INJ loading to 50% axial strain took 0.5s, Fig. 3C), and compressive (logarithmic) axial strain (damage initiation threshold: 40%^48,56,60^, Fig. 3D). All these mechanisms resulted in a substantial decrease in whole-thickness aggrecan content via aggrecanases over 12 days. The simulated aggrecan content was similar between the maximum shear strain-driven (coordinate system-independent strain measure) and fluid pressure-driven models, both estimating lower aggrecan content compared to the axial strain-driven model (coordinate system-dependent strain measure, Fig. 3E, left and middle panels). All three models were consistent with the experimental near/away from lesion aggrecan content fractions (day 7 experiment: 0.638±0.147; shear strain model 0.614; fluid pressure model 0.627; axial strain model 0.606, Fig. 3E, right panel).

**Fig. 3.**
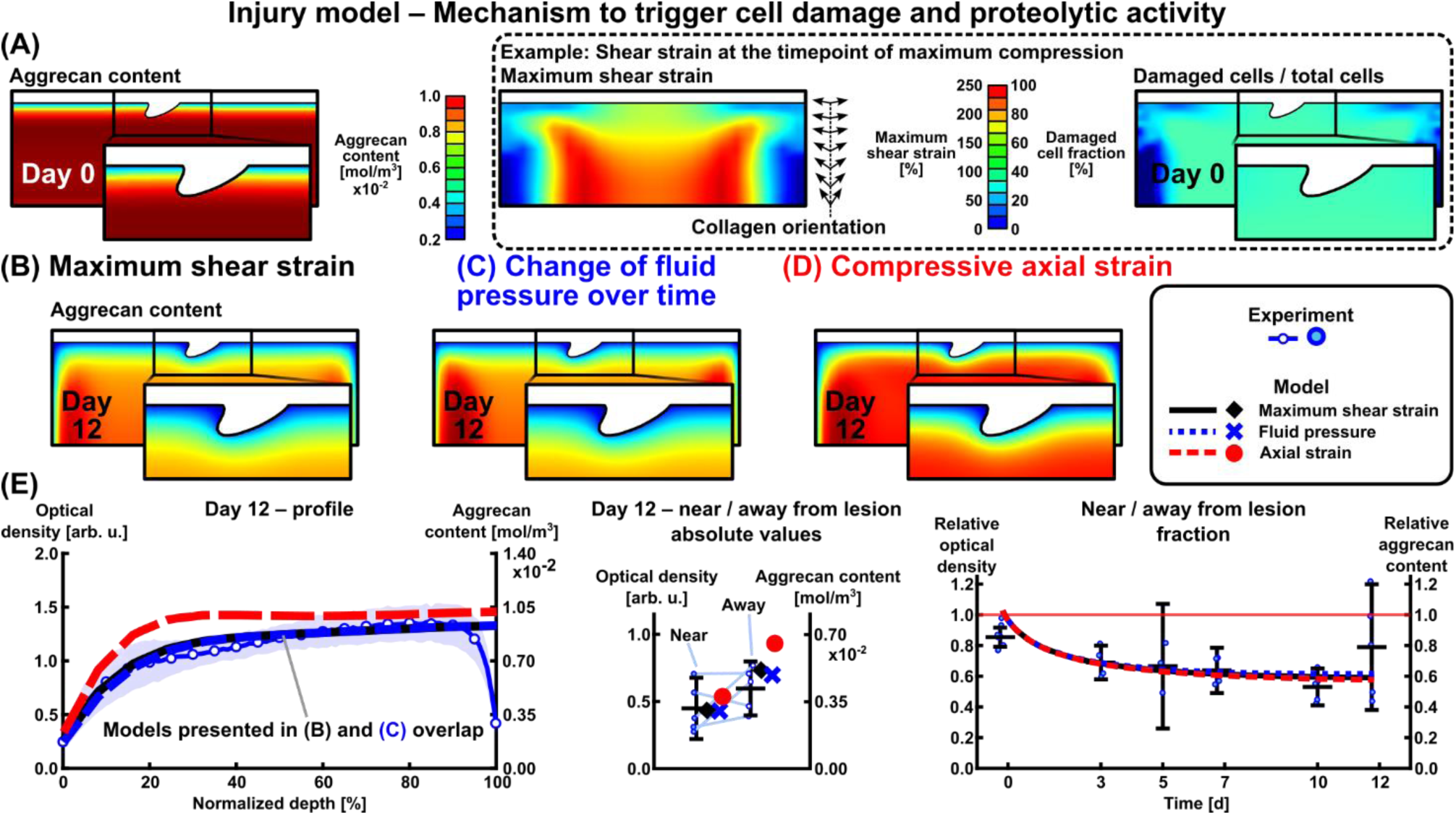
Different mechanisms to trigger cell damage and subsequent proteolytic activity (aggrecanase release) in the injury-only model. **(A, B)** Maximum shear strain (coordinate system-independent parameter) and **(C)** change of fluid pressure over time as mechanisms of cell damage resulted in similar estimates of aggrecan loss. **(D)** Compressive axial strain (coordinate system-dependent parameter) resulted in less aggrecan degradation than the maximum shear strain mechanism when all the model parameters were kept constant, as observed in **(E)** depth-wise, and near and away from lesion aggrecan contents. Maximum shear strain was selected for the rest of the study due to its coordinate system independence and more support from literature for the degradation thresholds as opposed to challenging-to-measure fluid pressures

Sensitivity analysis of the maximum shear strain-driven INJ model revealed that the threshold for shear strain-driven cellular damage *ε*_dmg,max_(in regions exceeding this threshold, fraction *k*_inj_ = 0.45 of healthy cells were considered damaged cells) had a minor effect on the absolute aggrecan content on day 12 (largest difference: model with *ε*_dmg,max_ = 200% resulted in 4.6% higher near-lesion aggrecan content than model with *ε*_dmg,max_ = 100%, Fig. 4A, middle panel). Increasing the values of parameters describing the amount of damaged cells *k*_inj_and aggrecanase release *k*_aga_decreased the aggrecan content considerably on day 12 (*k*_inj_ = 60% resulted in 33.5% lower near-lesion aggrecan content than *k*_inj_ = 20% , Fig. 4B; *k*_aga_ = 0.375 ⋅ 10^−21^ mol resulted in 26.0% lower near-lesion aggrecan content than *k*_aga_ = 0.125 ⋅ 10^−21^ mol, Fig. 4C). These three parameters (*ε*_dmg,max_, *k*_inj_, and *k*_aga_) had negligible effect on the relative near/away from lesion aggrecan content over time (maximum difference 3.2%-points in models with different *k*_aga_on day 12, Fig. 4C, right panel). Selecting a higher initial depth-wise aggrecan content^62^ compared to the current digital densitometry-based content^7^ resulted in higher relative near/away aggrecan content over time (0.789^62^ *vs.* 0.591^7^ on day 12, Fig. 4D, right panel).

**Fig. 4.**
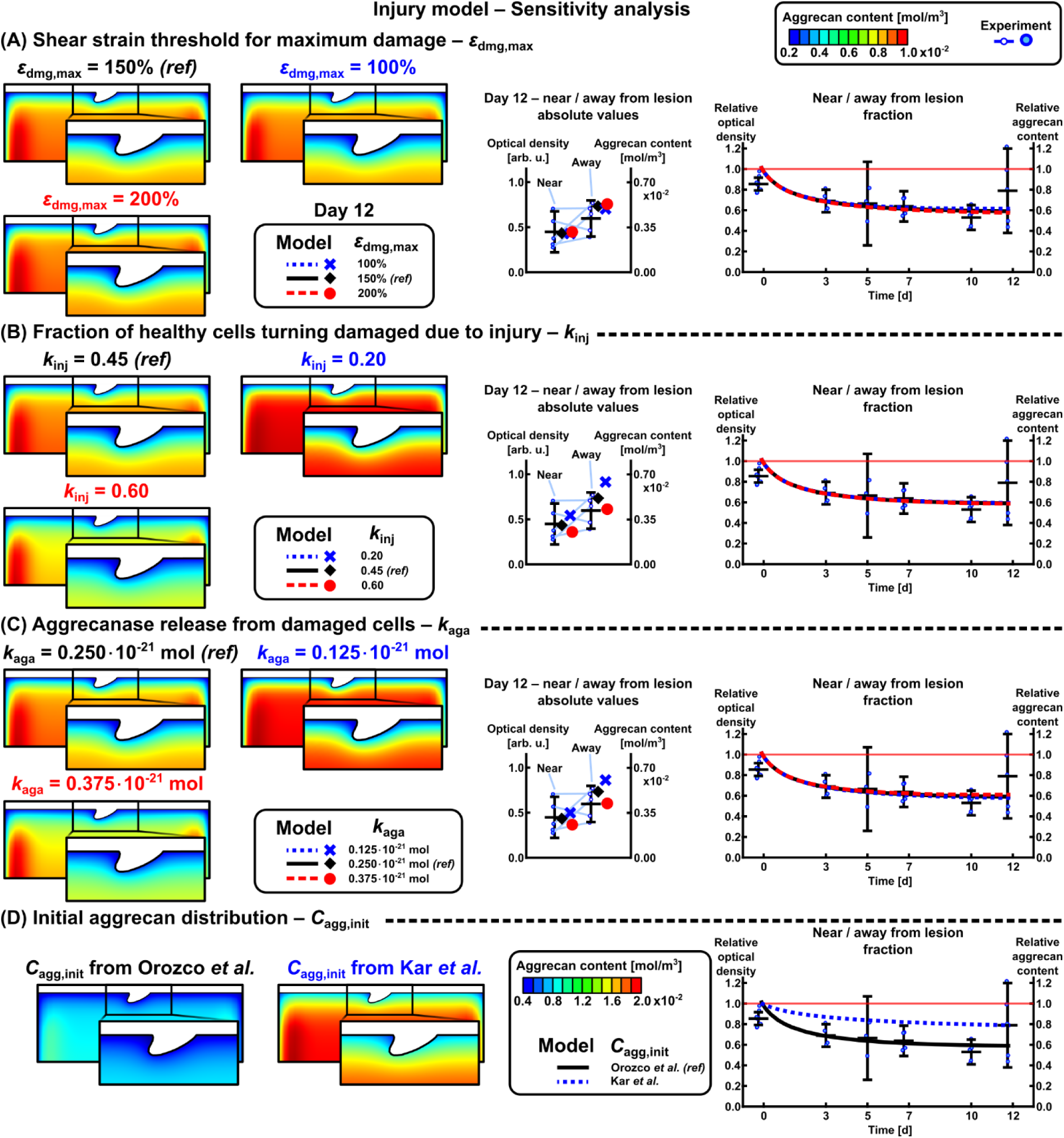
Effect of selected parameters on the aggrecan loss in the injurious loading model. **(A)** Maximum shear strain threshold value for maximum damage, **(B)** fraction of healthy cells turning damaged due to injury, and (**C)** proteolytic activity (aggrecanase release) scaled the absolute amounts of localized aggrecan content, but their effect on relative near *vs.* away aggrecan contents was negligible. On the other hand, **(D)** initial aggrecan concentration affected the relative change in aggrecan content near *vs.* away from lesions. Note different colorbar scale in (D); the current study uses Orozco *et al.*^7^ aggrecan distribution. *ref* = reference value selected to the models

### 2.4 Physiological loading promotes fluid flow-driven aggrecan loss near lesions and fluid pressure-stimulated acceleration of aggrecan biosynthesis in intact regions

The INJ+CL model was built step-by-step by first considering proteolytic activity only (localized cell damage from cyclic loading in addition to the initial injury-triggered catabolic cell activity, threshold of damage initiation 40%^48,56,60^, Fig. 5A). Due to mismatch with experimental near/away from lesion aggrecan content, additional cartilage adaptation mechanisms were investigated: elevated fluid flow velocity-related aggrecan depletion (threshold 0.08 mm ⋅ s^−1^ ^7,48^, Fig. 5B) and acceleration of aggrecan biosynthesis by up to 100% from basal biosynthesis rate in regions where fluid pressure changes were within a healthy range (20–60 MPa ⋅ s^−1^^22,59,63^ (the CL to 15% axial strain took 0.2s, Fig. 5C). Implementing the fluid flow mechanism increased near-lesion aggrecan loss compared to the model with proteolytic activity only (Fig. 5D, middle panel). Implementing the fluid pressure- stimulated acceleration of biosynthesis resulted in higher aggrecan content especially in away-from- lesion regions compared to models without accelerated biosynthesis (Fig. 5D, left and middle panels). On day 12, the relative near/away from lesion aggrecan content was 0.566 with proteolytic activity only, decreasing to 0.500 with proteolytic activity and fluid flow mechanisms combined, and to 0.452 with proteolytic activity, fluid flow, and fluid pressure-stimulated acceleration of aggrecan biosynthesis acting simultaneously (experiment: 0.411±0.053, Fig. 5D, right panel). Next, we used the model to numerically estimate the rate of aggrecan content increase/decrease associated to each of these mechanisms on day 12 (magnitude of source/sink terms in Eq. (4)). Near the lesion, the effect of fluid flow was 29.1% higher than that of proteolytic activity (Fig. 5E, left panel). Away from the lesion, the effect of fluid pressure-stimulated acceleration of aggrecan biosynthesis was 28.0% higher than the proteolytic effect (Fig. 5E, middle panel). Below the lesion and in deeper cartilage away from the lesion, the effect of accelerated biosynthesis was 22.7–27.2% lower than the proteolytic effect (Fig. 5E, right panel), with biosynthetic activity being lower below the lesion compared to away from the lesion.

**Fig. 5.**
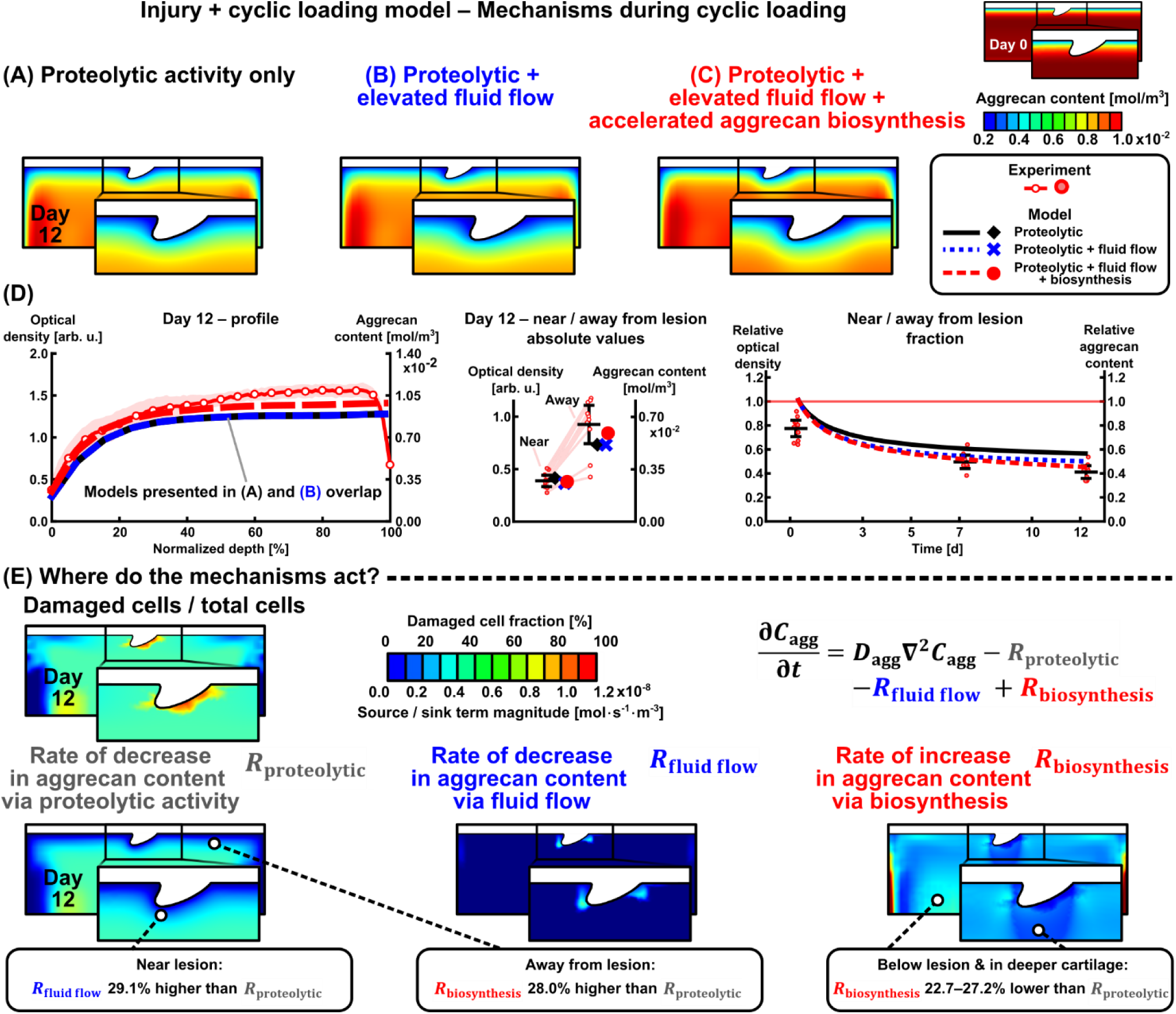
Different mechanisms associated with cyclic loading in the injury and cyclic loading model. **(A)** Only proteolytic activity due to damaged cells (triggered by maximum shear strain), **(B)** addition of the effect of elevated fluid flow flushing aggrecan fragments out of the tissue, and **(C)** addition of the effect of accelerated aggrecan biosynthesis when loading is within healthy limits (∼change of fluid pressure over time). **(D)** Depth-wise, and near and away from lesion aggrecan contents. **(E)** Cell-driven proteolytic activity affects the whole geometry, elevated fluid flow acts near the lesion, and accelerated aggrecan biosynthesis plays a role in the bulk of the model excluding near- and below-lesion regions

From sensitivity analysis of the INJ+CL model, we highlight that increasing the rate of cell damage *k*_cl_ and rate constant of aggrecan depletion due to fluid flow *k*_cl,fl_ decreased aggrecan content near lesion on day 12 (*k*_cl_ = 7.5 ⋅ 10^−6^ s^−1^ resulted in 5.4%-points lower near/away from lesion aggrecan content than *k*_cl_ = 0.3 ⋅ 10^−6^ s^−1^, Fig. 6A; *k*_cl,fl_ = 7.5 ⋅ 10^−6^ s^−1^ resulted in 16.8%-points lower near/away from lesion aggrecan content than *k*_cl,fl_ = 0.3 ⋅ 10^−6^ s^−1^, Fig. 6B). The threshold for accelerated aggrecan biosynthesis *p̂*_synth,init_ had negligible effect on near/away from lesion aggrecan content (maximum difference 3.2%-points, Fig. 6C), while lesion geometry affected by 10.9%-points on day 12 (confidence interval, Fig. 6D).

**Fig. 6.**
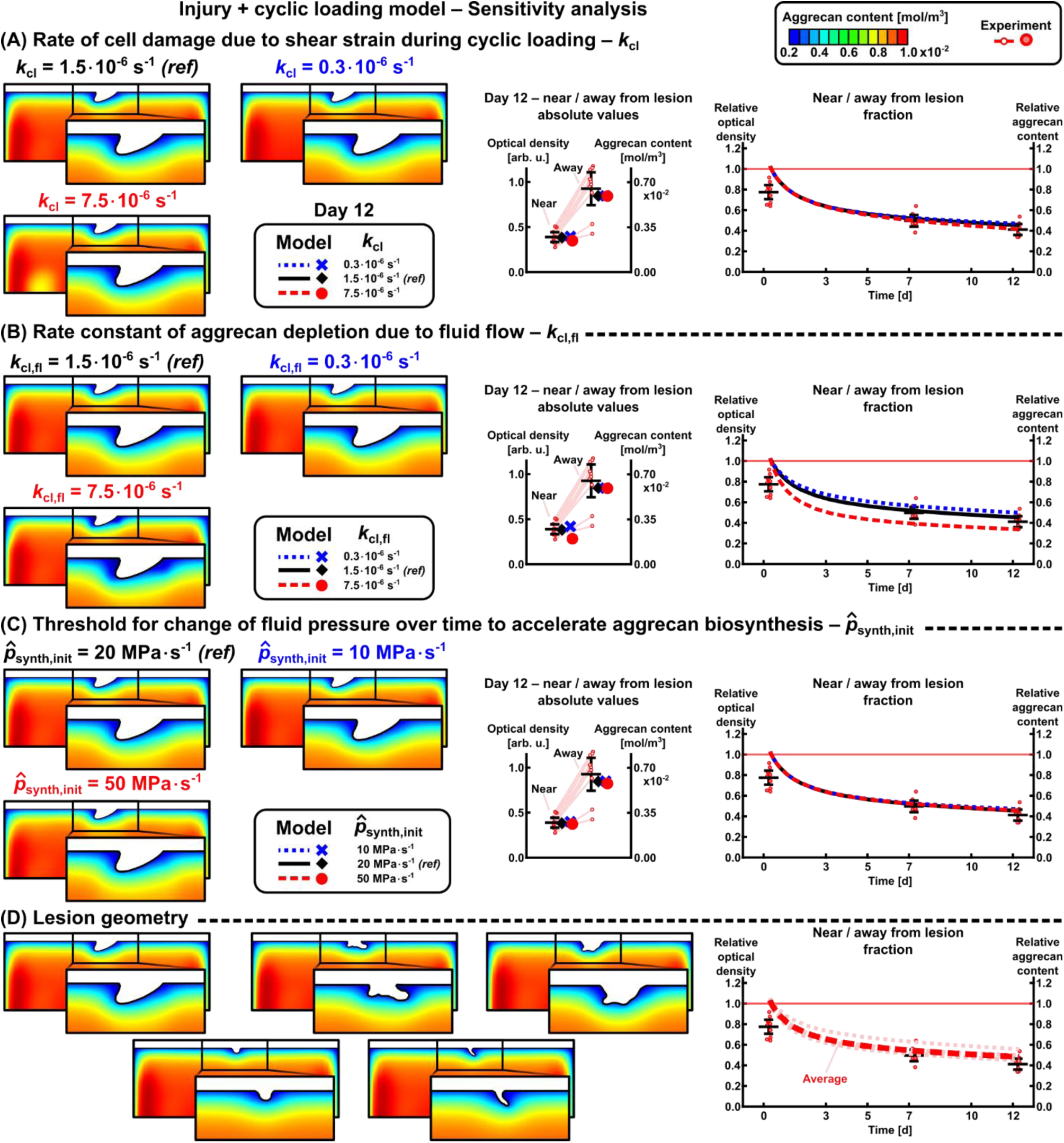
Effect of selected parameters on the aggrecan content adaptation in the injury and cyclic loading model. **(A)** Rate of cell damage (upregulating proteolytic activity) and **(B)** rate constant for aggrecan depletion due to fluid flow scaled the aggrecan loss especially near the lesion. **(C)** Fluid pressure-stimulated acceleration of aggrecan biosynthesis was present especially away from lesions, resulting in similar aggrecan content regardless of the chosen threshold. **(D)** Lesion geometry affected relative values of near vs. away from lesion aggrecan content for ∼25%-points on day 12. *ref* = reference value selected to the models

## 3 Discussion

We developed time-dependent computational models of cartilage mechanoinflammation to estimate changes in spatial aggrecan content under injurious and physiological cyclic loading. The models replicated experimental findings from two prior explant culture studies of early-stage PTOA^7,9^ in terms of depth-wise aggrecan loss (whole-thickness response), near and away from chondral lesion aggrecan loss (localized response), and near *vs.* away-from-lesion aggrecan content ratios (sample-specific difference in localized responses; Fig. 2). Implementing **(1)** proteolytic activity (aggrecanase release) from cells damaged due to excessive maximum shear strains (at day-0 injury and over the 12-day CL), **(2)** fluid flow flushing out aggrecan fragments, and **(3)** fluid pressure-stimulated acceleration of aggrecan biosynthesis in moderately loaded areas, resulted in ∼7% and ∼14% higher aggrecan loss near lesions with CL compared to without CL after 3 and 12 days from the injury, respectively (Fig. 2F). For those interested, the non-localized bulk aggrecan losses observed in these experimental setups were consistent with prior findings from the literature^10,14^, as discussed in detail elsewhere^7,9^. In line with our hypothesis, the mechanical response to CL in injured cartilage geometry was heterogeneous (depth-wise, near lesions). This resulted in the fluid flow effect dominating over proteolytic activity near the chondral lesions, accelerated aggrecan biosynthesis partially counteracting aggrecanase-driven aggrecan loss in the deep cartilage regions, and diminished biosynthetic activity below the lesions compared to regions away from them (Fig. 5E). Overall, this computational model offers a new basis for assessing biomechanics-driven spatio-temporal cartilage remodeling (both degradation and pro-anabolic responses). With further calibration, it could be utilized to design various rehabilitation exercises or tissue-engineered constructs to tackle PTOA progression.

Simulated injurious loading resulted in cell damage throughout the depth of the explant (Fig. 3A, right panel). This depth-wise cellular response aligns with reports of elevated oxidative stress and apoptosis/necrosis within the top 1 mm of injured cartilage^13,14,28,32^. We combined these cell fates into a “damaged cell phenotype” population; the fraction of damaged to total cells after injury (∼30– 45%, Fig. 3A) fell in the range of experimental studies with unconfined compression (20–50%^5^, 40– 55%^16^) and impact loading injury (40–65%^32^). The cell damage from injurious loading was assumed to occur immediately on day 0, although the timescales for different cell damage mechanisms vary from seconds to ∼1 h to ∼2 days^13,16,64^. The strain threshold *ε*_dmg,init_ = 40% for initiating damage was based on computational studies^48,60^ and has been suggested to be lower (7–12%)^58,65^. Using a lower value would essentially result in a visually similar aggrecan distribution as when increasing aggrecanase production (Fig 4C). Instead, the threshold *ε*_dmg,max_scaling maximum damage had negligible effect on aggrecan content (Fig. 4A).

Localized release of aggrecanase from damaged cells was one of the key assumptions of the model. This cellular response can be detected locally with ADAMTS immunostaining^66^, a potential method for finetuning the proteolytic parameter *k*_aga_ that was found to strongly affect aggrecan content (Fig. 4C). We based the value of *k*_aga_ on the work of Kar *et al.*^62^ — which focused on pro-inflammatory cytokine-driven aggrecan loss induced by interleukin (IL)-1 — with an assumption that stimulus for aggrecanase release would be similar for injurious loading (50% strain in 0.5s) and 1 ng/ml of IL-1^9,14^. As a technical sidenote, *k*_aga_*C*_cell,damaged_(Eq. S15) was in a similar range as the equilibrium concentration of IL-1–IL-1-receptor complexes *C*\* in^62^. A major part of immediate aggrecan loss is due to the microdamage from injury itself^4,67^, which is not accounted for in the model, rendering the value of *k*_aga_ an overestimate of the underlying damage mechanism.

Besides maximum shear strain (a coordinate system-independent strain measure), we also observed that coordinate system-dependent axial strain (Fig. 3D) and fluid pressure change over time (Fig. 3C) could be used to trigger cell damage and replicate the experimental findings. The axial strain is usable in simple geometries, while the shear strain and fluid pressure could serve as viable computational biomarkers of cell damage in more complex joint-level models^68^. However, caution must be taken with the suggested and challenging-to-validate fluid pressure damage threshold (80 MPa ⋅ s^−1^) serving as an upper limit where beneficial effects of loading turn detrimental^10,14,22^.

Simulated 12-day CL on injured cartilage resulted in progressive and localized cell damage^64^ but, as expected, without noticeable bulk damage as with INJ (Figs. 3A, 5E). The associated proteolytic activity from CL alone did not capture the observed aggrecan loss (Fig. 5D, right panel), necessitating the addition of the fluid flow mechanism to which aggrecan loss was sensitive near lesions (Figs. 5BE, 6B). The selected fluid flow velocity threshold with reasonable model fit (0.08 mm ⋅ s^−1^) was higher than the previously suggested^7^ value of 0.04 mm ⋅ s^−1^. However, the results from this modeling framework (same shear strain threshold in both INJ and CL, aggrecanase concentration set to zero on boundaries^62^) suggest that fluid flow dominates over proteolytic activity in the depletion of aggrecan fragments near lesions during CL (Fig. 5E), although this remains challenging to confirm experimentally.

Pro-anabolic effects of CL were added as accelerated aggrecan biosynthesis to improve model fitting to experimental aggrecan loss in intact and deep regions (Fig. 5D, left and middle panels). Increasing the basal aggrecan biosynthesis rate only in healthy cells by up to 100% (66–95%^69^, 20– 120%^40^; ^35^S-sulfate incorporation assays after similar CL protocols) partially counteracted the depth- wise proteolytic activity in deep and intact regions (Fig. 5E, right panel), with aggrecan content showing low sensitivity to the suggested biosynthesis thresholds (10–50 MPa ⋅ s^−1^ , Fig. 6C). Aggrecan biosynthesis was relatively low below the central lesion (Fig. 5E), consistent with reports of decreasing hydrostatic pressure gradient, biosynthesis rate, and availability of solutes/nutrients from central to lateral regions during unconfined compression^3,70^. Alterations in aggrecan biosynthesis could also be attributed to the activation of latent growth factors^3^ and localized energy dissipation to the tissue^71^, although these were not modeled here.

A limitation of this computational model is the inclusion of only two cell populations, “healthy” and “damaged”. Cartilage mechanoinflammation is far more complex, involving *e.g.* mechanical activation of ion channels (Piezo1/2, transient receptor potential vanilloid TRPV4) and downstream phenomena such as oxidative stress, mitochondrial dysfunction, apoptosis/necrosis, and cell hypertrophy^13,18,30,32,72^. The affected cell populations may orchestrate time-dependent reparative/remodeling responses to injury such as upregulating production of ADAMTS and tissue inhibitors of matrix metalloproteinases in surrounding viable cells (healthy and/or damaged) and, depending on the microenvironment, switching damaged cells to dead or back to healthy phenotype^4^. We chose to focus on replicating experimental findings with a minimal number of cell crosstalk- related parameters, which are explored elsewhere^3,49,53–55^. The model also omits the tissue responses to injury-related release of pro-inflammatory cytokines^14,50^ as well as lesion formation/propagation^73^. Thus, this study provides an estimate of the upper limit for the parameters of mechanical loading- related cell responses (peak biomechanical response used as input here instead of time-averaged estimates of the whole 12 days) and ECM remodeling in a single experimental setup. While utilizing young bovine cartilage provides a repeatable setup in terms of mechanical and biological tissue properties, it must be acknowledged that the present numerical estimates of mechanoinflammatory processes may not be fully translatable to more mature cartilage. Moreover, the results are dependent on the experimental protocol; use of different loading amplitudes, frequencies, and larger sample sizes could improve the robustness of the model parameters across different loading scenarios.

Degradation softens cartilage, potentially leading to locally increasing strains over time. This has been captured in previous models^7,51^ using an iterative approach, where ABAQUS material model was updated in each iteration (arbitrary unit of time, necessitating the introduction of new parameters). During model development, we noticed a ∼5% increase in maximum shear strains near the lesion in CL model with decreased aggrecan content (day 12 content of INJ+CL model), which would reduce aggrecan content approximately by <5% (Eq. (1), *k*_cl_in Fig. 6A). While tissue softening may not be a significant factor over 12 days, it becomes important over longer times in joint-level models.

In conclusion, by implementing several mechanoinflammatory cartilage adaptation mechanisms, the current modeling framework offers an avenue for studying the interplay between different loading scenarios, cartilage degradation and, as a relatively uncommon feature in the contemporary finite element modeling literature of osteoarthritis, the pro-anabolic responses in geometries with experimentally observed chondral lesions. The degradative and beneficial aspects of loading as presented here can be extended to knee joint level models to develop subject-specific osteoarthritis progression models^51^ and evaluate the effects of physical rehabilitation schemes^68,74^ to study cartilage damage for example after sports injuries. More generally, the modeling framework could also be developed further to simulate anti-catabolic and pro-anabolic therapeutic strategies^75^. Thus, the present study provides one piece to the multi-scale computational approaches aimed at helping limit PTOA progression.

## 4 Methods

### 4.1 Experimental groups and timeline

The experiments have been described in detail in two previous studies which investigated localized aggrecan loss in early-stage PTOA^7,44^. This study utilizes a subset of those data. Briefly, cartilage explants (thickness 1 mm, diameter 3 mm, *n*=105) were harvested from patellofemoral grooves of 1– 2-week-old calves on the day of slaughter (*N*=10 animals, one knee per animal, Research 87 Inc., Boylston, MA). After equilibration for two days in serum-free culture medium (marking day 0), the explants were divided into the following groups: free-swelling control (CTRL, *n*=19, cultures terminated on days 0 and 12)^7^, injurious loading only (INJ, *n*=54, days 0^7,44^, 3, 5, 7, 10, and 12^44^), cyclic loading only (CL, *n*=11, day 12)^7^, and injury and cyclic loading groups (INJ+CL, *n*=21, days 7 and 12)^7^.

### 4.2 Mechanical loading protocols

On day 0, explants in the INJ and INJ+CL groups were subjected to a single load–unload cycle of unconfined compression with 50% axial strain relative to the sample-specific thickness (strain rate 100%/s) within a custom-designed incubator-housed loading apparatus^10,14,44,76,77^. The resulting peak stresses ranged from 20–30 MPa (peak force per top surface area of the undeformed explant). This protocol has been demonstrated to decrease cell viability, especially near lesions^5,7,14,44^, and to cause both bulk and localized loss of aggrecan content^8,14,44^.

The CL and INJ+CL explants were subjected to cyclic loading protocol designed to mimic physiological loading and daily activities. After establishing repeatable contact with the explants with 10% compressive offset strain (five 2% axial strain ramp-and-hold increments, each increment followed by a six-minute stress-relaxation), the explants were subjected continuously to cyclic unconfined compression with 15% axial strain (1 Hz, haversine waveform, 40% duty cycle, 1-h loading sessions four times a day) in an incubator-housed cyclic loading apparatus^7,14,44^. This CL protocol has previously been shown to be non-detrimental for the bulk cartilage (negligible aggrecan loss^7^, increased aggrecan biosynthesis^14,44^) with the potential to inhibit pro-catabolic responses in cartilage following mechanical injury^14^.

### 4.3 Analysis of aggrecan content

Aggrecan content was estimated by measuring optical density (arbitrary units) in three 3-μm-thick, Safranin-O-stained cartilage sections per explant using digital densitometry (conventional light microscope, calibration with series of filters of known optical density, Nikon Microphot FXA, Nikon Inc., Tokyo, Japan, 4× magnification, pixel size 1.23 μm)^44,78^. Injured explants with sections showing excessive tissue rupture or no lesions were excluded from the analysis (14/75 explants, 18.7%). For each section, optical density was measured in the following regions: **(1)** full-thickness depth-wise profiles (two 200-μm-wide regions, one from either side of lesion, averaged along the width and then depth-wise, Fig. 1B), **(2)** near lesions (average value within 50 μm from lesion edges; mean lesion depth 135±18 μm (95%CI), mean lesion width 204±45 μm), **(3)** away from lesions (same analysis region shape as near lesion, translated to both sides of lesion and averaged; mean distance from lesion edges 635±100 μm). For **(4)** relative near *vs*. away from lesion fractions, the average optical density value near lesion was divided by the average away-from-lesion value. Lastly, each measure was averaged over the three sections per explant.

### 4.4 Statistics

All statistical analyses were conducted using linear mixed effects (LME) models in IBM SPSS Statistics 29.0 (SPSS Inc., IBM Company, Armonk, NY, USA). Data points more than 1.5 times the interquartile range below the 25% quartile or above the 75% quartile were discarded as outliers. Different animals were considered as subjects with random intercepts, and treatment, time, and patellofemoral surface region where each explant was harvested from, as fixed factors (in the statistical analysis, the combined dataset had four surface regions from one study^44^ and one region from the other^7^). Location-matching was done by harvesting explants for all treatment groups from each patellofemoral region. The covariance structure was set as the variance components structure (default). The significance level was set to α = 0.05, and results are reported with Bonferroni correction.

### 4.5 Computational models of mechanical loading

Two unconfined compression loading models were developed in ABAQUS (v. 2023, Dassault Systèmes Simulia Corp.) according to the experimental protocols: one for the initial single injurious compression on day 0 (50% strain in 0.5s; 100%/s) and another for the physiological cyclic loading (15% strain, with 40% duty cycle the maximum strain was obtained in 0.2s, two cycles modeled^7^). With cartilage modeled as a fibril-reinforced porohyperelastic swelling material incorporating depth-dependent compositional properties^7,79^ (Supplementary Table S1), the injurious loading was subjected to intact geometry (peak stresses ∼50 MPa when using material parameters found from literature with which model was still converging, higher than ∼30 MPa in experiments). Lesion formation or crack propagation^73^ were not modeled. Cyclic loading was subjected to intact (CL group) or injured geometry (INJ+CL group, five representative lesions of varying sizes were created based on the cartilage sections, Fig. 6D). All bottom surface nodes were fixed in vertical direction and in addition the middle bottom node was fixed in the horizontal direction, allowing for lateral expansion of the tissue. Fluid flow was allowed through the lateral and lesion surfaces (pore pressure = 0). Mesh convergence for both the injurious loading (Supplementary Fig. S1) and cyclic loading models^7^ was assured. The models were developed in 2D to facilitate the comparison of simulated and experimentally observed aggrecan loss in cartilage sections with lesions.

The mechanical loading models provided input parameters to the time-dependent mechanoinflammatory cartilage adaptation models (see Section 4.6). These included maximum shear strain as a trigger for day-0 cellular initial conditions in INJ (during model development, also fluid pressure change over time and logarithmic axial strain, Fig. 3), and maximum shear strain, fluid flow, and fluid pressure change over time (promoting accelerated aggrecan biosynthesis) in the 12-day simulations of CL and INJ+CL (Fig. 5, Supplementary Material S1). The parameters were obtained at the timepoint of maximum compression; the parameter ‘fluid pressure change over time’ (rate of pressure buildup has been suggested to affect cellular responses^4^) was calculated between the beginning of the compression and the maximum compression (in ABAQUS, pore pressure change between two time points divided by the time period; over 0.5s in INJ, over 0.2s in CL and INJ+CL).

### 4.6 Computational models of mechanoinflammatory cartilage adaptation

The biomechanical model outputs from ABAQUS (such as, maximum shear strain *ε*) served as inputs for the mechanoinflammatory cartilage adaptation models with spatio-temporally evolving aggrecan concentration (COMSOL Multiphysics, v. 5.6., COMSOL AB, Stockholm, Sweden). The injurious loading was assumed to trigger initial cell damage on day 0, implemented with a normalized shear strain-driven cellular damage function *f*_dmg_(*ε*)^27,80^ as

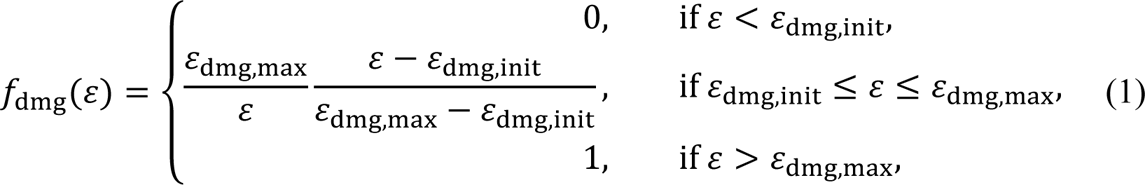

where thresholds for cell damage initiation *ε*_dmg,init_ = 40% ^48,56,60^ and for maximum damage *ε*_dmg,max_ = 150%^56^. Equations of this form were also used in the INJ models to investigate other potential cell damage triggers (excessive fluid pressure change over time and logarithmic compressive axial strain, Fig. 3CD, Table S2). The damage function *f*_dmg_(*ε*) was used to convert part of the initial concentration of healthy cells *C*_cell,healthy,init_ into damaged cells *C*_cell,damaged,init_ on the day of injury^15,56^ with the following initial condition^55^:

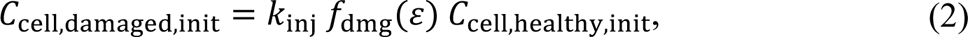

where *k*_inj_ = 0.45 is the maximum fraction of healthy cells turning to damaged phenotype in young bovine cartilage after injurious compression/impact^5,16,32,54^. Due to the lack of suitable quantitative data, this modeling approach which involved only two cell populations aimed to represent the overall phenotypic switching from “healthy” to “damaged” cells without explicitly considering the underlying cell damage mechanisms separately (apoptosis^5,15,54^, necrosis^15,54,64^, oxidative stress^13,54,69^, mitochondrial dysfunction^81^, hypertrophic state^3,82^). Depending on cellular microenvironment, the damage may be considered minor (altered cellular function) or major (cell death), with both scenarios associated with aggrecanase release either from the damaged cells themselves or from viable cells adjacent to damaged regions^4^. Therefore, as a generalization, we assumed that the modeled entity “damaged cells” would release aggrecanases (ADAMTS-4,5, a disintegrin and metalloproteinase with thrombospondin motifs) that diffuse within the cartilage ECM and eventually decrease the aggrecan content^54,66^. The cartilage adaptation was modeled with time-dependent diffusion–reaction equations over the 12 days following injury^50,62^:

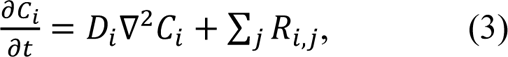

where *C_i_* is the time-dependent concentration of the species *i* (healthy cells, damaged cells, aggrecan, aggrecanase), *D_i_* is effective diffusivity (0 for both cell populations), *t* is time, and *R_i,j_* are the source (+)/sink (−) terms for species *i* and mechanism *j* (*e.g.*, aggrecanase activity, aggrecan biosynthesis). For instance, the aggrecan concentration *C*_agg_ evolved as:

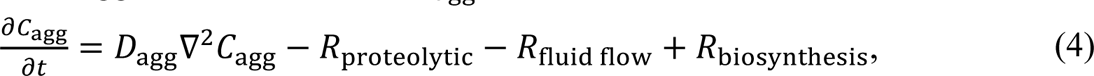

where the proteolytic activity decreasing aggrecan content *R*_proteolytic_ depended on the release of aggrecanases from damaged cells (catabolic cell activity driven by both injury on day 0 and cyclic loading over 12 days; for the full equations about the cascade of mechanical stimulus – mechanoinflammatory response – aggrecanase release – aggrecan degradation, see Supplementary Material S1, Table S2). Elevated fluid flow-driven aggrecan depletion *R*_fluid flow_ depended on the magnitude of fluid flow velocity^7,48,83^, and the addition of new aggrecan via biosynthesis *R*_biosynthesis_depended on the change of fluid pressure over time in areas with healthy cells^10^ experiencing fluid pressure alterations in the range considered healthy (20–60 MPa ⋅ s^−1^ over 0.2s of cyclic loading^22,59,61,63^). The maximum increase in the aggrecan biosynthesis rate was set to 100%^40,44,69^.

Due to limited experimental evidence for the model parameter values, a sensitivity analysis was conducted to assess how the most relevant model parameters affected the estimates of aggrecan content. The reference parameters (Table 1) represent the INJ and INJ+CL models, which predicted the average aggrecan contents based on experimental data. The parameter value ranges were selected based on the literature and/or by adjusting the reference value to obtain a reasonable fit to the upper/lower bounds of the confidence intervals. In addition, two different initial aggrecan concentrations^7,62^ and five different lesion geometries were investigated.

**Table 1.**
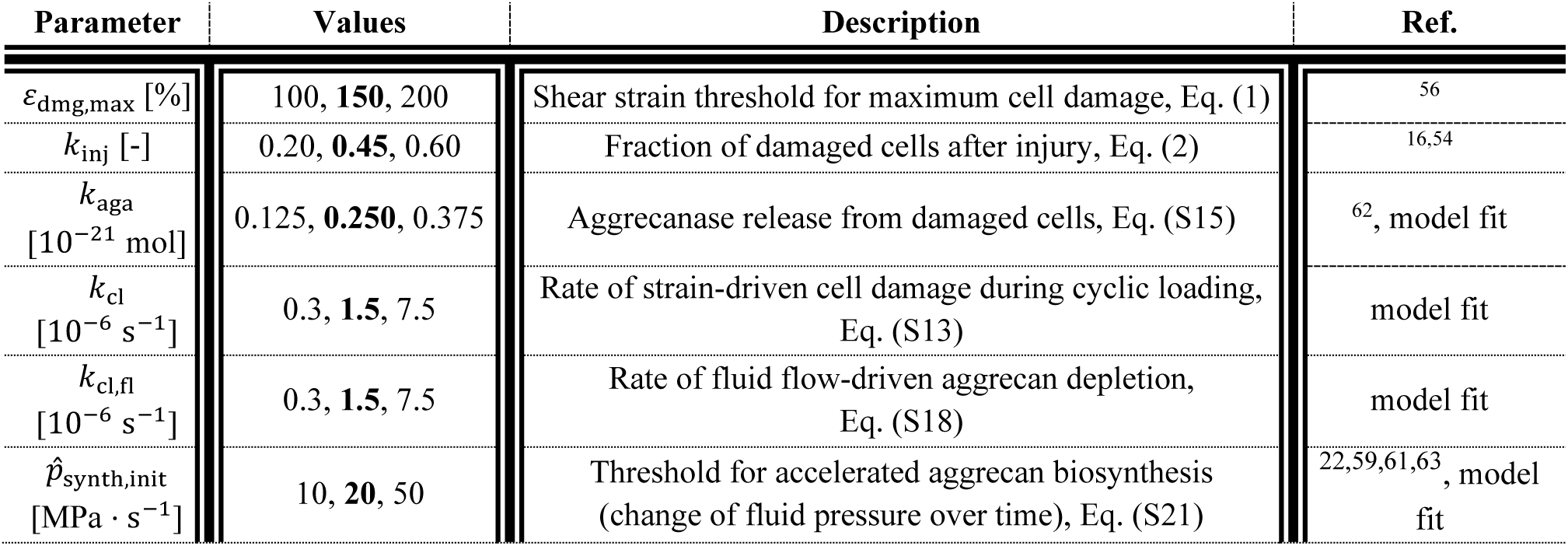
Parameters selected to the sensitivity analysis. Bolded values represent reference values.

## Data availability

The datasets generated and analyzed during this study are available from the corresponding author upon reasonable request.

## Acknowledgements

This work was supported by the Doctoral Programme in Science, Technology and Computing (LUMETO) of the University of Eastern Finland; the Research Council of Finland (grant numbers 354916, 363459); Strategic Funding of the University of Eastern Finland; the Sigrid Jusélius Foundation; the Instrumentarium Science Foundation; the Saastamoinen Foundation; the Finnish Cultural Foundation (grants 191044 and 00230091); the Maire Lisko Foundation; the Novo Nordisk Foundation (grant NNF21OC0065373); This project has received funding from the European Union’s Horizon 2020 research and innovation programme under the Marie Skłodowska-Curie grant agreement Nos. 702586 and 101108335. The funding sources were not involved in the data collection, analysis, or interpretation. Eija Rahunen (Institute of Biomedicine, Cell and Tissue Imaging Unit, UEF) is acknowledged for the processing of the samples designated for digital densitometry analysis.

## Author contributions

**Atte Eskelinen:** Conceptualization, Methodology, Formal analysis, Investigation, Data curation, Writing – Original Draft, Writing – Review & Editing, Visualization. **Joonas Kosonen:** Conceptualization, Methodology, Writing – Review & Editing, Visualization. **Moustafa Hamada:** Conceptualization, Methodology, Writing – Review & Editing. **Amir Esrafilian:** Conceptualization, Methodology, Writing – Review & Editing. **Cristina Florea**: Conceptualization, Methodology, Writing – Review & Editing, Visualization, Supervision. **Petri Tanska:** Conceptualization, Methodology, Writing – Review & Editing, Visualization, Supervision. **Alan Grodzinsky:** Conceptualization, Methodology, Resources, Writing – Review & Editing, Supervision, Project administration, Funding acquisition. **Rami Korhonen**: Conceptualization, Methodology, Resources, Writing – Review & Editing, Visualization, Supervision, Project administration, Funding acquisition.

## Supporting information

**S1 Supplementary Material.** Electronic supplementary material containing more detailed information.

**Fig. S1. Mesh sensitivity analysis.**

**Table S1. Material parameters in the mechanical loading model.**

**Table S2. Model parameters in the mechanoinflammatory cartilage adaptation model.**

## References

1. Mobasheri, A. et al. Recent advances in understanding the phenotypes of osteoarthritis. F1000Res 8, 1–11 (2019).

2. Deveza, L. et al. Knee osteoarthritis phenotypes and their relevance for outcomes: a systematic review. Osteoarthritis Cartilage 25, 1926– 1941 (2017).

3. Rahman, M. M., Watton, P. N., Neu, C. P. & Pierce, D. M. A chemo-mechano-biological modeling framework for cartilage evolving in health, disease, injury, and treatment. Comput Methods Programs Biomed 231, (2023).

4. Riegger, J. & Brenner, R. E. Pathomechanisms of posttraumatic osteoarthritis: Chondrocyte behavior and fate in a precarious environment. Int J Mol Sci 21, (2020).

5. Patwari, P. et al. Ultrastructural quantification of cell death after injurious compression of bovine calf articular cartilage. Osteoarthritis Cartilage 12, 245–252 (2004).

6. Hodgkinson, T., Kelly, D. C., Curtin, C. M. & O’Brien, F. J. Mechanosignalling in cartilage: an emerging target for the treatment of osteoarthritis. Nat Rev Rheumatol 18, 67–84 (2022).

7. Orozco, G., Tanska, P., Florea, C., Grodzinsky, A. & Korhonen, R. A novel mechanobiological model can predict how physiologically relevant dynamic loading causes proteoglycan loss in mechanically injured articular cartilage. Sci Rep 8, 15599 (2018).

8. Patwari, P., Cheng, D. M., Cole, A. A., Kuettner, K. E. & Grodzinsky, A. J. Analysis of the relationship between peak stress and proteoglycan loss following injurious compression of human post-mortem knee and ankle cartilage. Biomech Model Mechanobiol 6, 83–89 (2007).

9. Eskelinen, A. S. A. et al. Cyclic loading regime considered beneficial does no protect injured and interleukin-1-inflamed cartilage from post-traumatic osteoarhtritis. 1–50 (2021).

10. Kurz, B. et al. Biosynthetic response and mechanical properties of articular cartilage after injurious compression. Journal of Orthopaedic Research 19, 1140–1146 (2001).

11. Hines, M. R. et al. Extracellular biomolecular free radical formation during injury. Free Radic Biol Med 188, 175–184 (2022).

12. Coleman, M. C. et al. Targeting mitochondrial responses to intra-articular fracture to prevent posttraumatic osteoarthritis. Sci Transl Med 10, 1–32 (2018).

13. Coleman, M. C., Ramakrishnan, P. S., Brouillette, M. J. & Martin, J. A. Injurious Loading of Articular Cartilage Compromises Chondrocyte Respiratory Function. Arthritis and Rheumatology 68, 662–671 (2016).

14. Li, Y. et al. Moderate Dynamic Compression Inhibits Pro-Catabolic Response of Cartilage to Mechanical Injury, TNF-α and IL-6, but Accentuates Degradation Above a Strain Threshold. Osteoarthritis Cartilage 21, (2013).

15. Stolberg-Stolberg, J. A. et al. Effects of cartilage impact with and without fracture on chondrocyte viability and the release of inflammatory markers. Journal of Orthopaedic Research 31, 1283–1292 (2013).

16. Loening, A. M. et al. Injurious mechanical compression of bovine articular cartilage induces chondrocyte apoptosis. Arch Biochem Biophys 381, 205–212 (2000).

17. Martin, J. A., McCabe, D., Walter, M., Buckwalter, J. A. & McKinley, T. O. N-acetylcysteine inhibits post-impact chondrocyte death in osteochondral explants. Journal of Bone and Joint Surgery - Series A 91, 1890–1897 (2009).

18. Bartell, L., Fortier, L., Bonassar, L. & Cohen, I. Measuring microscale strain fields in articular cartilage during rapid impact reveals thresholds for chondrocyte death and a protective role for the superficial layer. J Biomech 48, 3440–3446 (2015).

19. DiMicco, M. A., et al. Mechanisms and Kinetics of Glycosaminoglycan Release Following In Vitro Cartilage Injury. Arthritis Rheum 50, 840–848 (2004).

20. Larsson, T., Aspden, R. M. & Heinegård, D. Effects of mechanical load on cartilage matrix biosynthesis in vitro. Matrix 11, 388–394 (1991).

21. Rieder, B. et al. Hydrostatic pressure-generated reactive oxygen species induce osteoarthritic conditions in cartilage pellet cultures. Sci Rep 8, 1–16 (2018).

22. Hall, A. C., Urban, J. P. G. & Gehl, K. A. The effects of hydrostatic pressure on matrix synthesis in articular cartilage. Journal of Orthopaedic Research 9, 1–10 (1991).

23. Occhetta, P. et al. Hyperphysiological compression of articular cartilage induces an osteoarthritic phenotype in a cartilage-on-a-chip model. Nat Biomed Eng 3, 545–557 (2019).

24. Chen, C.-T., Burton-Wurster, N., Lust, G., Bank, R. A. & Iekoppele, J. M. Compositional and metabolic changes in damaged cartilage are peak-stress, stress-rate, and loading-duration dependent. Journal of Orthopaedic Research 17, 870–879 (1999).

25. Komeili, A., Abusara, Z., Federico, S. & Herzog, W. Effect of strain rate on transient local strain variations in articular cartilage. J Mech Behav Biomed Mater 95, 60–66 (2019).

26. Párraga Quiroga, J. M., Wilson, W., Ito, K. & van Donkelaar, C. C. Relative contribution of articular cartilage’s constitutive components to load support depending on strain rate. Biomech Model Mechanobiol 16, 151–158 (2017).

27. Párraga Quiroga, J., Wilson, W., Ito, K. & van Donkelaar, C. The effect of loading rate on the development of early damage in articular cartilage. Biomech Model Mechanobiol 16, 263–273 (2017).

28. Morel, V. & Quinn, T. M. Cartilage injury by ramp compression near the gel diffusion rate. Journal of Orthopaedic Research 22, 145–151 (2004).

29. Singh, P., Marcu, K. B., Goldring, M. B. & Otero, M. Phenotypic instability of chondrocytes in osteoarthritis: On a path to hypertrophy. Ann N Y Acad Sci 1–18 (2018) doi:10.1111/nyas.13930.

30. Henak, C. R., Bartell, L. R., Cohen, I. & Bonassar, L. J. Multiscale Strain as a Predictor of Impact-Induced Fissuring in Articular Cartilage. J Biomech Eng 139, 031004 (2017).

31. Delco, M. L., Bonnevie, E. D., Bonassar, L. J. & Fortier, L. A. Mitochondrial dysfunction is an acute response of articular chondrocytes to mechanical injury. Journal of Orthopaedic Research 36, 739–750 (2018).

32. Ayala, S., Delco, M. L., Fortier, L. A., Cohen, I. & Bonassar, L. J. Cartilage articulation exacerbates chondrocyte damage and death after impact injury. Journal of Orthopaedic Research 39, 2130–2140 (2021).

33. Lee, J. H. et al. Co-culture of mechanically injured cartilage with joint capsule tissue alters chondrocyte expression patterns and increases ADAMTS5 production. Arch Biochem Biophys 489, 118–126 (2009).

34. Verma, P. & Dalal, K. ADAMTS-4 and ADAMTS-5: Key enzymes in osteoarthritis. J Cell Biochem 112, 3507–3514 (2011).

35. Larsson, S., Lohmander, L. S. & Struglics, A. Biological variation of human aggrecan ARGS neoepitope in synovial fluid and serum in early-stage knee osteoarthritis and after knee injury. Osteoarthr Cartil Open 4, 100307 (2022).

36. Didomenico, C. D., Lintz, M. & Bonassar, L. J. Molecular transport in articular cartilage - What have we learned from the past 50 years? Nat Rev Rheumatol 14, 393–403 (2018).

37. Soltz, M. A. & Ateshian, G. A. Interstitial Fluid Pressurization During Confined Compression Cyclical Loading of Articular Cartilage. (2000).

38. Sharma, G., Saxena, R. K. & Mishra, P. Differential effects of cyclic and static pressure on biochemical and morphological properties of chondrocytes from articular cartilage. Clinical Biomechanics 22, 248–255 (2007).

39. Sah, R. et al. Biosynthetic response of cartilage explants to dynamic compression. Journal of Orthopaedic Research 7, 619–636 (1989).

40. Kisiday, J. D., Jin, M., DiMicco, M. A., Kurz, B. & Grodzinsky, A. J. Effects of dynamic compressive loading on chondrocyte biosynthesis in self-assembling peptide scaffolds. J Biomech 37, 595–604 (2004).

41. Kisiday, J. D. et al. Catabolic responses of chondrocyte-seeded peptide hydrogel to dynamic compression. Ann Biomed Eng 37, 1368–1375 (2009).

42. DiMicco, M. A. & Sah, R. L. Dependence of cartilage matrix composition on biosynthesis, diffusion, and reaction. Transp Porous Media 50, 57–73 (2003).

43. Säämänen, A. -M, Tammi, M., Kiviranta, I., Jurvelin, J. & Helminen, H. J. Levels of chondroitin-6-sulfate and nonaggregating proteoglycans at articular cartilage contact sites in the knees of young dogs subjected to moderate running exercise. Arthritis Rheum 32, 1282–1292 (1989).

44. Eskelinen, ASA. et al. Cyclic loading regime considered beneficial does not protect injured and interleukin-1-inflamed cartilage from post-traumatic osteoarthritis. J Biomech 141, 111181 (2022).

45. Venäläinen, M. et al. Quantitative evaluation of the mechanical risks caused by focal cartilage defects in the knee. Sci Rep 6, 37538 (2016).

46. Zevenbergen, L. et al. Functional assessment of strains around a full-thickness and critical sized articular cartilage defect under compressive loading using MRI. Osteoarthritis Cartilage 26, 1710–1721 (2018).

47. Zevenbergen, L. et al. Cartilage defect location and stiffness predispose the tibiofemoral joint to aberrant loading conditions during stance phase of gait. PLoS One 13, 1–22 (2018).

48. Orozco, G. A. et al. Prediction of local fixed charge density loss in cartilage following ACL injury and reconstruction: A computational proof-of-concept study with MRI follow-up. Journal of Orthopaedic Research 39, 1–8 (2020).

49. Kapitanov, G. I., Ayati, B. P. & Martin, J. A. Modeling the effect of blunt impact on mitochondrial function in cartilage: Implications for development of osteoarthritis. PeerJ 2017, 1–18 (2017).

50. Eskelinen, A. S. A. et al. Mechanobiological model for simulation of injured cartilage degradation via proinflammatory cytokines and mechanical stimulus. PLoS Comput Biol 16, 1–25 (2020).

51. Mononen, M. E., Tanska, P., Isaksson, H. & Korhonen, R. K. New algorithm for simulation of proteoglycan loss and collagen degeneration in the knee joint: Data from the osteoarthritis initiative. Journal of Orthopaedic Research 36, 1673–1683 (2018).

52. Moo, E. K., Ebrahimi, M., Sibole, S. C., Tanska, P. & Korhonen, R. K. The intrinsic quality of proteoglycans, but not collagen fibres, degrades in osteoarthritic cartilage. Acta Biomater (2022) doi:10.1016/j.actbio.2022.09.002.

53. Gardiner, B. et al. Predicting knee osteoarthritis. Ann Biomed Eng 44, 222–233 (2016).

54. Kosonen, J. P. et al. Injury-related cell death and proteoglycan loss in articular cartilage: Numerical model combining necrosis, reactive oxygen species, and inflammatory cytokines. PLoS Comput Biol 19, e1010337 (2023).

55. Kapitanov, G., Wang, X., Ayati, B., Brouillette, M. & Martin, J. Linking cellular and mechanical processes in articular cartilage lesion formation: A mathematical model. Front Bioeng Biotechnol 4, 1–22 (2016).

56. Argote, P. F. et al. Chondrocyte viability is lost during high-rate impact loading by transfer of amplified strain, but not stress, to pericellular and cellular regions. Osteoarthritis Cartilage 27, 1822–1830 (2019).

57. Huang, W., Warner, M., Sasaki, H., Furukawa, K. S. & Ushida, T. Layer dependence in strain distribution and chondrocyte damage in porcine articular cartilage exposed to excessive compressive stress loading. J Mech Behav Biomed Mater 112, (2020).

58. Bonnevie, E. et al. Microscale frictional strains determine chondrocyte fate in loaded cartilage. J Biomech 74, 72–78 (2018).

59. Elder, B. D. & Athanasiou, K. A. Hydrostatic Pressure in Articular Cartilage Tissue Engineering: From Chondrocytes to Tissue Regeneration. Tissue Engineering: Part B 15, 1–12 (2009).

60. Orozco, G. A. et al. Shear strain and inflammation-induced fixed charge density loss in the knee joint cartilage following ACL injury and reconstruction: a computational study. Journal of Orthopaedic Research 40, 1505–1522 (2022).

61. Bachrach, N. M., Mow, V. C. & Guilak, F. Incompressibility of the solid matrix of articular cartilage under high hydrostatic pressures. J Biomech 31, 445–451 (1998).

62. Kar, S. et al. Modeling IL-1 induced degradation of articular cartilage. Arch Biochem Biophys 594, 37–53 (2016).

63. Morrell, K. C., Andrew Hodge, W., Krebs, D. E. & Mann, R. W. Corroboration of in Vivo Cartilage Pressures with Implications for Synovial Joint Tribology and Osteoarthritis Causation. www.pnas.orgcgidoi10.1073pnas.0507117102 (2005).

64. Chen, C. T. et al. Chondrocyte necrosis and apoptosis in impact damaged articular cartilage. Journal of Orthopaedic Research 19, 703–711 (2001).

65. Ayala, S. et al. Degradation of lubricating molecules in synovial fluid alters chondrocyte sensitivity to shear strain. Journal of Orthopaedic Research (2024) doi:10.1002/jor.25960.

66. Merrild, N. G. et al. Local depletion of proteoglycans mediates cartilage tissue repair in an ex vivo integration model. Acta Biomater 149, 179–188 (2022).

67. Quinn, T. M., Maung, A. A., Grodzinsky, A. J., Hunziker, E. B. & Sandy, J. D. Physical and biological regulation of proteoglycan turnover around chondrocytes in cartilage explants. Implications for tissue degradation and repair. Ann N Y Acad Sci 878, 420–441 (1999).

68. Esrafilian, A. et al. Effects of gait modifications on tissue-level knee mechanics in individuals with medial tibiofemoral osteoarthritis: A proof-of-concept study towards personalized interventions. Journal of Orthopaedic Research (2023) doi:10.1002/jor.25686.

69. Momin, A., Perrotti, S. & Waldman, S. D. The role of mitochondrial reactive oxygen species in chondrocyte mechanotransduction. Journal of Orthopaedic Research (2023) doi:10.1002/jor.25709.

70. Kim, Y.-J., Sah, R. L. Y., Grodzinsky, A. J., Plaas, A. H. K. & Sandy, J. D. Mechanical Regulation of Cartilage Biosynthetic Behavior: Physical Stimuli. Arch Biochem Biophys 311, 1–12 (1994).

71. Stampoultzis, T., Guo, Y., Nasrollahzadeh, N., Karami, P. & Pioletti, D. P. Mimicking Loading-Induced Cartilage Self-Heating in Vitro Promotes Matrix Formation in Chondrocyte-Laden Constructs with Different Mechanical Properties. ACS Biomater Sci Eng 9, 651–661 (2023).

72. Zhang, M., Meng, N., Wang, X., Chen, W. & Zhang, Q. TRPV4 and PIEZO channels mediate the mechanosensing of chondrocytes to the biomechanical microenvironment. Membranes (Basel) 12, 1–11 (2022).

73. Orozco, G. A., Tanska, P., Gustafsson, A., Korhonen, R. K. & Isaksson, H. Crack propagation in articular cartilage under cyclic loading using cohesive finite element modeling. J Mech Behav Biomed Mater 131, (2022).

74. Esrafilian, A. et al. Toward Tailored Rehabilitation by Implementation of a Novel Musculoskeletal Finite Element Analysis Pipeline. IEEE Transactions on Neural Systems and Rehabilitation Engineering 30, 789–802 (2022).

75. Black, R. & Grodzinsky, A. J. Dexamethasone: Chondroprotective corticosteroid or catabolic killer? Eur Cell Mater 38, 246–263 (2019).

76. Frank, E. H., Jin, M., Loening, A. M., Levenston, M. E. & Grodzinsky, A. J. A versatile shear and compression apparatus for mechanical stimulation of tissue culture explants. J Biomech 33, 1523–1527 (2000).

77. Patwari, P. et al. Proteoglycan degradation after injurious compression of bovine and human articular cartilage in vitro: Interaction with exogenous cytokines. Arthritis Rheum 48, 1292–1301 (2003).

78. Király, K. et al. Application of selected cationic dyes for the semiquantitative estimation of glycosaminoglycans in histological sections of articular cartilage by microspectrophotometry. Histochemical Journal 28, 577–590 (1996).

79. Wilson, W., van Donkelaar, C., van Rietbergen, B. & Huiskes, R. A fibril-reinforced poroviscoelastic swelling model for articular cartilage. J Biomech 38, 1195–1204 (2005).

80. Hosseini, S., Wilson, W., Ito, K. & van Donkelaar, C. A numerical model to study mechanically induced initiation and progression of damage in articular cartilage. Osteoarthritis Cartilage 22, 95–103 (2014).

81. Brouillette, M. J. et al. Strain-dependent oxidant release in articular cartilage originates from mitochondria. Biomech Model Mechanobiol 13, 565–572 (2014).

82. Rim, Y. A., Nam, Y. & Ju, J. H. The role of chondrocyte hypertrophy and senescence in osteoarthritis initiation and progression. Int J Mol Sci 21, 2358 (2020).

83. Mow, V. C., Wang, C. C. & Hung, C. T. The extracellular matrix, interstitial fluid and ions as a mechanical signal transducer in articular cartilage. Osteoarthritis Cartilage 7, 41–58 (1999).

