## Supplementary Material S1 for "Time-dependent computational model of post-traumatic osteoarthritis to estimate how mechanoinflammatory mechanisms impact cartilage aggrecan content"

### Abbreviations

INJ = injurious loading

CL = cyclic loading

### S1 Biomechanical model parameters (ABAQUS)

Cartilage was modeled as a fibril-reinforced porohyperelastic material with Donnan osmotic swelling[1,2]. For the non-fibrillar part (proteoglycans), the Cauchy stress tensor of a neo-Hookean solid material is

$$\boldsymbol{\sigma}_{\mathrm{nf}} = K_{\mathrm{nf}} \frac{\ln(J)}{J} \mathbf{I} + \frac{G_{\mathrm{nf}}}{J} \left( \mathbf{F} \cdot \mathbf{F}^{\mathrm{T}} - J^{\frac{2}{3}} \mathbf{I} \right), \qquad \text{(Eq. S1)}$$

where $\mathbf{F}$ is the deformation gradient tensor, $J$ = det($\mathbf{F}$) is volumetric deformation, and $\mathbf{I}$ is the unit tensor. $K_{\mathrm{nf}}$ and $G_{\mathrm{nf}}$ are the bulk and shear moduli of the non-fibrillar matrix, respectively,

$$K_{\mathrm{nf}} = \frac{E_{\mathrm{nf}}}{3(1 - 2\nu_{\mathrm{nf}})}, \qquad \text{(Eq. S2)}$$

$$G_{\mathrm{nf}} = \frac{E_{\mathrm{nf}}}{2(1 + \nu_{\mathrm{nf}})}, \qquad \text{(Eq. S3)}$$

where $E_{\mathrm{nf}}$ and $\nu_{\mathrm{nf}}$ are the Young´s modulus and Poisson´s ratio of the non-fibrillar matrix, respectively. The stress in a single collagen fibril (after approximating that viscoelastic damping coefficient $\eta \approx 0$) was modeled with a linear elastic spring (initial fibril network modulus $E_0$)

$$\sigma_{\mathrm{f}} = \begin{cases} E_0 \varepsilon_{\mathrm{f}}, & \text{if } \varepsilon_{\mathrm{f}} \geq 0, \\ 0, & \text{if } \varepsilon_{\mathrm{f}} < 0, \end{cases} \qquad \text{(Eq. S4)}$$

where $\varepsilon_{\mathrm{f}}$ is the logarithmic fibril strain ($\varepsilon_{\mathrm{f}}$ = ln(||$\mathbf{F}\boldsymbol{e}_{\mathrm{f},0}$||, where $\boldsymbol{e}_{\mathrm{f},0}$ is the unit vector of the initial fibril orientation). For a fibril $j$ the Cauchy stress tensor is

$$\boldsymbol{\sigma}_{\mathrm{f}}{}^{j} = \begin{cases} \rho_{\mathrm{z}} C \sigma_{\mathrm{f}} \boldsymbol{e}_{\mathrm{f}} \otimes \boldsymbol{e}_{\mathrm{f}}, & \text{for primary fibrils,} \\ \rho_{\mathrm{z}} \sigma_{\mathrm{f}} \boldsymbol{e}_{\mathrm{f}} \otimes \boldsymbol{e}_{\mathrm{f}}, & \text{for secondary fibrils,} \end{cases} \qquad \text{(Eq. S5)}$$

where $\rho_{\mathrm{z}}$ is the depth-dependent collagen fraction per solid volume, $C$ is the density ratio between primary and secondary fibrils and $\otimes$ symbolizes dyadic product. The collagen network consisted of 2 primary fibrils defining curved basic structure found in young calf cartilage and 7 secondary fibrils describing cross-links and randomly oriented fibrils[2]. $\boldsymbol{e}_{\mathrm{f}}$ is the current normalized fibril orientation vector

$$\boldsymbol{e}_{\mathrm{f}} = \frac{\mathbf{F}\boldsymbol{e}_{\mathrm{f},0}}{||\mathbf{F}\boldsymbol{e}_{\mathrm{f},0}||}. \qquad \text{(Eq. S6)}$$

The fluid flow was incorporated according to Darcy's law

$$q = -k\nabla p, \quad \text{(Eq. S7)}$$

where $q$ is the flow rate in the non-fibrillar matrix, $k$ is the hydraulic permeability (constant, since the strain-dependent permeability factor $M = 0$ in FRPHES model) and $\nabla p$ is the pressure gradient.

Donnan osmotic swelling pressure gradient in equilibrium is

$$\Delta\pi = \phi_{\mathrm{int}} RT\left(\sqrt{c_{\mathrm{F}}^2 + 4\frac{\left(\gamma_{\mathrm{ext}}^{\pm}\right)^2}{\left(\gamma_{\mathrm{int}}^{\pm}\right)^2} c_{\mathrm{ext}}^2}\right) - 2\phi_{\mathrm{ext}} RT c_{\mathrm{ext}}, \quad \text{(Eq. S8)}$$

where $c_{\mathrm{F}}$ is the current depth-dependent fixed charge density (FCD) concentration, $\phi_{\mathrm{int}}$, $\phi_{\mathrm{ext}}$, $\gamma_{\mathrm{int}}^{\pm}$ and $\gamma_{\mathrm{ext}}^{\pm}$ are internal and external osmotic coefficients and internal and external activity coefficients, respectively, $c_{\mathrm{ext}}$ is the external salt concentration (0.15 M), $R$ is the molar gas constant (8.314 J/mol K) and $T$ is the absolute temperature (293.0 K). The chemical expansion stress is

$$T_{\mathrm{c}} = a_0 c_{\mathrm{F}} \exp\left(-\kappa \frac{\gamma_{\mathrm{ext}}^{\pm}}{\gamma_{\mathrm{int}}^{\pm}} \sqrt{c^-\left(c^- + c_{\mathrm{F}}\right)}\right), \quad \text{(Eq. S9)}$$

where $a_0$ and $\kappa$ are material constants and $c^-$ is the mobile anion concentration. The current depth-dependent FCD concentration is modeled as a function of volumetric deformation

$$c_{\mathrm{F}} = c_{\mathrm{F},0} \frac{n_{\mathrm{f},0}}{n_{\mathrm{f},0} - 1 + J}, \quad \text{(Eq. S10)}$$

where $c_{\mathrm{F},0}$ is the initial depth-dependent FCD and $n_{\mathrm{f},0}$ is the initial fluid volume fraction (*i.e.*, porosity). The total stress tensor is

$$\boldsymbol{\sigma}_{\mathrm{tot}} = \sum_{j=1}^{totf} \boldsymbol{\sigma}_{\mathrm{f}}{}^{j} + \boldsymbol{\sigma}_{\mathrm{nf}} - \Delta\pi\mathbf{I} - T_{\mathrm{c}}\mathbf{I} - \mu_{\mathrm{f}}\mathbf{I}, \quad \text{(Eq. S11)}$$

where *totf* is the sum of primary and secondary fibrils (2 + 7 = 9) and $\mu_{\mathrm{f}}$ is the chemical potential of water. Maximum shear strain $\varepsilon$ was calculated in MATLAB as

$$\varepsilon = \max\left\{\left|\varepsilon_{\mathrm{p},1} - \varepsilon_{\mathrm{p},2}\right|, \left|\varepsilon_{\mathrm{p},1} - \varepsilon_{\mathrm{p},3}\right|, \left|\varepsilon_{\mathrm{p},2} - \varepsilon_{\mathrm{p},3}\right|\right\}, \quad \text{(Eq. S12)}$$

where $\varepsilon_{\mathrm{p},k}$ are the principal strains of the Green–Lagrangian strain tensor (deformation gradient tensor obtained from ABAQUS). Logarithmic axial strain was obtained from ABAQUS logarithmic strain components "LE" (see also Eq. (2), Fig. 3D). The material parameters are listed in Table S1.

**Table S1. Material model parameters (ABAQUS).** $z$ indicates normalized distance from the cartilage surface (surface = 0, bottom = 1) in the depth-dependent properties. The parameters represent experimental explants where the top 1 mm was cut from the biopsy-punched cartilage samples.

| Parameter | Value | Description | Reference |
|---|---|---|---|
| **Compositional** | | | |
| $n_{\mathrm{f},0}$ [-] | $0.85 - 0.1z$ | Initial fluid fraction in equilibrium | [1–4] |
| $\rho_{\mathrm{z}}$ [-] | $20.6z^6 - 64.4z^5 + 78.1z^4$ $-45.9z^3 + 13.4z^2 - 1.6z + 0.96$ | Collagen fraction | [2,4,5] |
| $c_{\mathrm{F},0}$ $[\mathrm{mEq} \cdot \mathrm{ml}^{-1}]$ | $-4.4z^6 + 15.2z^5 - 21.0z^4 +$ $14.9z^3 - 5.8z^2 + 1.1z + 0.03$ | Initial fixed charge density | [2,4,6] |
| **Material** | | | |
| $C$ [-] | 3.009 | Density ratio between primary and secondary collagen fibrils | [1,2,4] |
| $E_0$ [MPa] | 20 | Initial fibril network modulus | [2,4] |
| $E_{\mathrm{nf}}$ [MPa] | 0.16 | Non-fibrillar matrix modulus | [2,4] |
| $\nu_{\mathrm{nf}}$ [-] | 0.4 | Non-fibrillar matrix Poisson's ratio | [2,4,7] |
| $k$ $[10^{-15}\,\mathrm{m}^4 \cdot \mathrm{N}^{-1} \cdot \mathrm{s}^{-1}]$ | 1.3 | Hydraulic permeability | [2,4] |

#### S2 Mesh sensitivity analysis

Mesh sensitivity analysis was undertaken for the model of single injurious compression to ensure mesh-independence of the estimated biomechanical responses (maximum shear strain; Fig. S1). A similar analysis for the cyclic loading model has previously been done elsewhere[2,4]. The mesh with 799 elements was selected to provide injury-related initial conditions for the INJ and INJ+CL models as refining the mesh from there resulted in <2% change in maximum shear strain estimates.

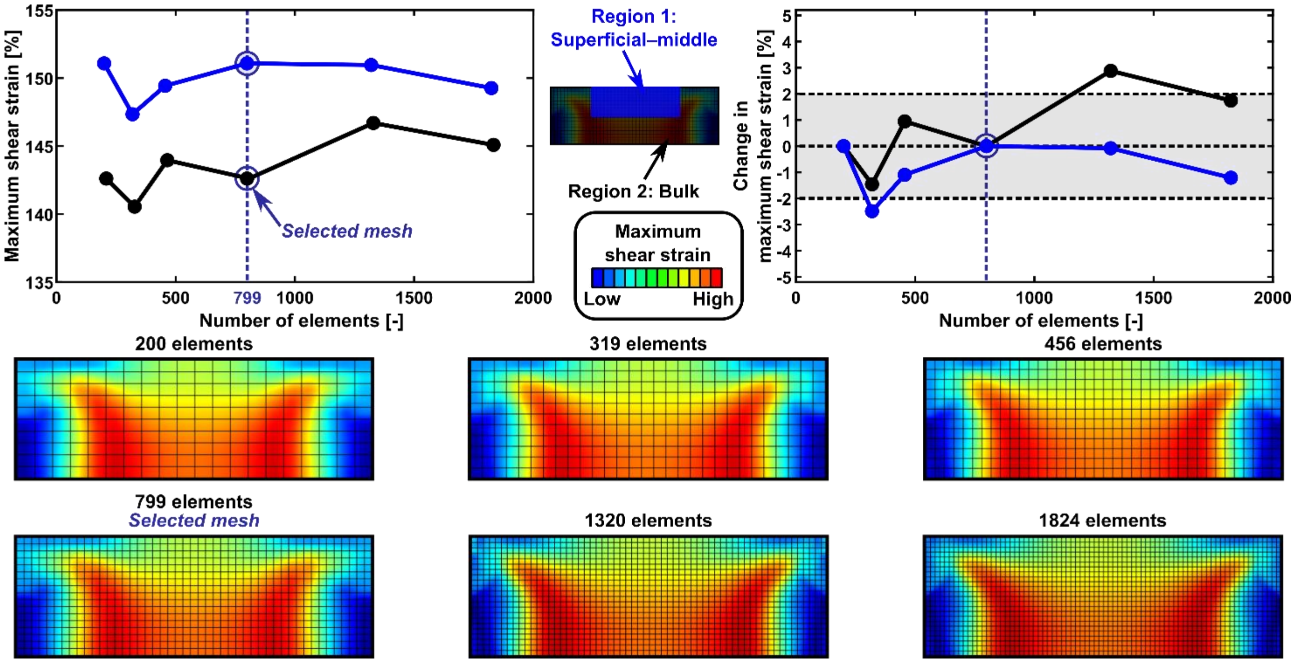


**Fig. S1. Mesh sensitivity analysis.** Different meshes for the injurious loading model were tested to ascertain that mesh is not influencing the results. The mechanical parameter triggering cell damage (maximum shear strain) is shown as an average in the superficial–middle area where lesion was formed (blue; width 1.5 mm, depth 0.5 mm) and bulk (black) at the time of maximum compression (50% axial strain, figure shows undeformed mesh). The mesh with 799 elements was selected for the study; its shear strain estimates deviated <2% from more refined meshes.

#### S3 Mechanoinflammatory cartilage adaptation model parameters (COMSOL Multiphysics)

Elevated maximum shear strains (Eq. S12) were hypothesized to turn healthy cells into damaged during injury (Eqs. 1 and 2) and during cyclic loading. In the latter, the normalized shear strain-driven cellular damage function $f_{\mathrm{dmg}}(\varepsilon)$ was calculated at the time of largest deformation during compressive cyclic loading. Healthy and damaged cell populations evolved over time as

$$\frac{\partial C_{\mathrm{cell,healthy}}}{\partial t} = -k_{\mathrm{cl}}\, f_{\mathrm{dmg}}(\varepsilon)\, C_{\mathrm{cell,healthy}}, \qquad \text{(Eq. S13)}$$

$$\frac{\partial C_{\mathrm{cell,damaged}}}{\partial t} = k_{\mathrm{cl}}\, f_{\mathrm{dmg}}(\varepsilon)\, C_{\mathrm{cell,healthy}}, \qquad \text{(Eq. S14)}$$

respectively, where $k_{\mathrm{cl}}$ is the rate of cell damage (rate of cells turning from healthy to damaged) due to cyclic loading. Concentration of damaged cells (both due to INJ and CL) gave rise to localized time-dependent stimulus for aggrecanase release; this (delayed) mechanoinflammatory stimulus term approach was motivated by Kar *et al.*[8] and findings that cell-driven enzymatic activity is delayed (in contrast to very early aggrecan loss due to microdamage from the mechanical insult) and increasing on the subsequent hours–days from injury[9] (delay due to trauma-related release of alarmins and damage-associated molecular patterns peaking in ~24h[10]).

$$\frac{\partial S_{\mathrm{aga}}}{\partial t} = \alpha_{\mathrm{aga}}\left(k_{\mathrm{aga}} C_{\mathrm{cell,damaged}} - S_{\mathrm{aga}}\right), \qquad \text{(Eq. S15)}$$

where $S_{\mathrm{aga}}$ is stimulus for aggrecanase, is $\alpha_{\mathrm{aga}}$ rate constant for aggrecanase stimulus, and $k_{\mathrm{aga}}$ is constant for aggrecanase release from damaged cells (calibrated so that the stimulus variables are in similar range than those in the adaptation models of Kar *et al.*[8]). The aggrecanase concentration $C_{\mathrm{aga}}$ evolved as

$$\frac{\partial C_{\mathrm{aga}}}{\partial t} = D^{*}_{\mathrm{aga}} e^{-d_1 C_{\mathrm{agg}}} \nabla^2 C_{\mathrm{aga}} + k_1 S_{\mathrm{aga}} - k_2 C_{\mathrm{aga}}, \qquad \text{(Eq. S16)}$$

where $D^{*}_{\mathrm{aga}}$ is diffusivity of aggrecanase, $d_1$ is a constant defining the dependency of aggrecanases' effective diffusivity on local aggrecan concentration ($D_{\mathrm{aga}} = D^{*}_{\mathrm{aga}} e^{-d_1 C_{\mathrm{agg}}}$), $k_1$ is rate constant for generating aggrecanases based on damaged cell-driven stimulus, and $k_2$ is enzymatic loss of aggrecanases. The aggrecan concentration $C_{\mathrm{agg}}$ was defined in Eq. (4), where the source/sink terms are

$$R_{\mathrm{proteolytic}} = k_3 C_{\mathrm{aga}} \frac{C_{\mathrm{agg}}}{C_{\mathrm{aga}} + K_{\mathrm{m,aga}}}, \qquad \text{(Eq. S17)}$$

where $k_3$ is catalytic rate constant for aggrecanase and $K_{\mathrm{m,aga}}$ is the Michaelis constant for aggrecanase, and

$$R_{\mathrm{fluid\ flow}} = k_{\mathrm{cl,fl}}\, f_{\mathrm{dmg}}(v)\, C_{\mathrm{agg}}, \qquad \text{(Eq. S18)}$$

where $k_{\mathrm{cl,fl}}$ is rate constant for aggrecan depletion due to elevated fluid flow and $f_{\mathrm{dmg}}(v)$ is normalized fluid flow velocity-driven matrix damage function, defined in the same manner as for maximum shear strain in Eq. (1):

$$f_{\mathrm{dmg}}(v) = \begin{cases} 0, & \text{if } v < v_{\mathrm{dmg,init}}, \\ \dfrac{v_{\mathrm{dmg,max}}}{v} \dfrac{v - v_{\mathrm{dmg,init}}}{v_{\mathrm{dmg,max}} - v_{\mathrm{dmg,init}}}, & \text{if } v_{\mathrm{dmg,init}} \le v \le v_{\mathrm{dmg,max}}, \\ 1, & \text{if } v > v_{\mathrm{dmg,max}}, \end{cases} \qquad \text{(Eq. S19)}$$

where $v$ is the magnitude of fluid flow velocity and $v_{\mathrm{dmg,init}}$ and $v_{\mathrm{dmg,max}}$ are thresholds for damage initiation and maximum non-fibrillar matrix damage, respectively. The aggrecan biosynthesis term in the aggrecan diffusion–reaction equation is

$$R_{\mathrm{biosynthesis}} = \left(1 + f_{\mathrm{synth}}(\hat{p})\right) P_{\mathrm{ag}} \left(1 + 0.9\frac{1 - z}{H}\right) C_{\mathrm{cell,healthy}} \left(1 - \frac{C_{\mathrm{agg}}}{C_{\mathrm{agg,tar}}}\right), \qquad \text{(Eq. S20)}$$

where $P_{\mathrm{ag}}$ is chondrocyte-based basal aggrecan production (biosynthesis rate), $z$ is normalized depth in cartilage ($z = 0$ surface, $z = 1$ bottom), $H = 1$ mm is cartilage thickness, and $C_{\mathrm{agg,tar}}$ is target (homeostatic) aggrecan concentration. The normalized biosynthesis function $f_{\mathrm{synth}}(\hat{p})$ could upregulate aggrecan biosynthesis rate $P_{\mathrm{ag}}$ locally by up to 100%[11,12], defined as

$$f_{\mathrm{synth}}(\hat{p}) = \begin{cases} 0, & \text{if } \hat{p} < \hat{p}_{\mathrm{synth,init}}, \\ \dfrac{\hat{p}_{\mathrm{synth,max}}}{\hat{p}} \dfrac{\hat{p} - \hat{p}_{\mathrm{synth,init}}}{\hat{p}_{\mathrm{synth,max}} - \hat{p}_{\mathrm{synth,init}}}, & \text{if } \hat{p}_{\mathrm{synth,init}} \leq \hat{p} \leq \hat{p}_{\mathrm{synth,max}}, \\ 1, & \text{if } \hat{p} > \hat{p}_{\mathrm{synth,max}}, \end{cases} \quad \text{(Eq. S21)}$$

where $\hat{p}$ is the change of fluid (pore) pressure over time ($\mathrm{MPa \cdot s^{-1}}$; between beginning of compression and the time point of largest deformation, that is, at 15% strain after 0.2 s in the CL and INJ+CL models), and $\hat{p}_{\mathrm{synth,init}}$ and $\hat{p}_{\mathrm{synth,max}}$ are the thresholds ($\mathrm{MPa \cdot s^{-1}}$) for initiation and maximum upregulation of aggrecan biosynthesis rate. Moreover, similar equation was used to trigger cell damage with excessive fluid pressure change over time on the day of injury ($\hat{p}_{\mathrm{dmg,init}}$ and $\hat{p}_{\mathrm{dmg,max}}$ for cell damage initiation and maximum damage, respectively, see also Eq. (2), Fig. 3C). The parameter values are listed in Table S2.

**Table S2. Adaptation model distributions, boundary conditions, and parameters (COMSOL Multiphysics).**

| Parameter | Value / Expression | Description | Reference |
|---|---|---|---|
| **Distributions** | | | |
| $C_{\mathrm{cell,healthy,init}}$ $[\mathrm{cells \cdot m^{-3}}]$ | $1.5 \cdot 10^{14}$ | Initial healthy cell distribution | 4,6,8 |
| $C_{\mathrm{agg,init}}$ $[\mathrm{mol \cdot m^{-3}}]$ | $\dfrac{c_{\mathrm{F,0}} \cdot 502.5}{-2 \cdot 2.5 \cdot 1000}$ | Initial aggrecan distribution, derived from fixed charge density (Table S1) | 2,13 |
| **Boundary conditions** | | | |
| | $D_{\mathrm{agg}} \nabla C_{\mathrm{agg}} = 0$ | Aggrecan flux through bottom | 4,6,8 |
| | $D_{\mathrm{agg}} \nabla C_{\mathrm{agg}} + h_{\mathrm{z,agg}}(C_{\mathrm{agg}} - C_{\mathrm{agg,b}}) = 0$ | Aggrecan flux through top | 4,6,8 |
| | $D_{\mathrm{agg}} \nabla C_{\mathrm{agg}} + h_{\mathrm{r,agg}}(C_{\mathrm{agg}} - C_{\mathrm{agg,b}}) = 0$ | Aggrecan flux through lateral edges | 4,6,8 |
| | $D_{\mathrm{aga}} \nabla C_{\mathrm{aga}} = 0$ | Aggrecanase flux through bottom | 4,6,8 |
| | $C_{\mathrm{aga}} = 0$ | Aggrecanase concentration at top and lateral edges | 4,6,8 |
| **Parameters** | | | |
| $\varepsilon_{\mathrm{dmg,init}}$ [%] | 40 | Shear/logarithmic axial threshold for cell damage initiation | 13–15 |

| | | | |
|---|---|---|---|
| $\varepsilon_{\mathrm{dmg,max}}$ [%] | 150 | Shear/logarithmic axial strain strain threshold for maximum cell damage | [14] |
| $v_{\mathrm{dmg,init}}$ $[\mathrm{mm} \cdot \mathrm{s}^{-1}]$ | 0.08 | Fluid flow velocity threshold for non-fibrillar matrix damage initiation | [13,15], model fit |
| $v_{\mathrm{dmg,max}}$ $[\mathrm{mm} \cdot \mathrm{s}^{-1}]$ | 0.15 | Fluid flow velocity threshold for maximum non-fibrillar matrix damage | [13,15] |
| $\hat{p}_{\mathrm{dmg,init}}$ $[\mathrm{MPa} \cdot \mathrm{s}^{-1}]$ | 80 | Threshold for initiating cell damage due to excessive pore pressure change over time | [16–18], model tests |
| $\hat{p}_{\mathrm{dmg,max}}$ $[\mathrm{MPa} \cdot \mathrm{s}^{-1}]$ | 100 | Threshold for maximum cell damage due to excessive pore pressure change over time | [16–18], model tests |
| $\hat{p}_{\mathrm{synth,init}}$ $[\mathrm{MPa} \cdot \mathrm{s}^{-1}]$ | 20 | Threshold for initiating acceleration of aggrecan biosynthesis rate due to moderate pore pressure change over time | [16–18], model tests |
| $\hat{p}_{\mathrm{synth,max}}$ $[\mathrm{MPa} \cdot \mathrm{s}^{-1}]$ | 60 | Threshold for maximum aggrecan biosynthesis rate due to moderate pore pressure change over time | [16–18], model tests |
| $k_{\mathrm{inj}}$ [-] | 0.45 | Maximum fraction of healthy cells turning damaged after injury | [6,19] |
| $D_{\mathrm{agg}}$ $[\mathrm{m}^2 \cdot \mathrm{s}^{-1}]$ | $10^{-14}$ | Effective diffusivity of aggrecan | [8] |
| $D^*_{\mathrm{aga}}$ $[\mathrm{m}^2 \cdot \mathrm{s}^{-1}]$ | $10^{-12}$ | Diffusivity of aggrecanase | [8] |
| $d_1$ $[\mathrm{m}^3 \cdot \mathrm{mol}^{-1}]$ | 120 | Constant defining the dependency of effective diffusivity of aggrecanase to aggrecan concentration | [8] |
| $k_{\mathrm{cl}}$ $[\mathrm{s}^{-1}]$ | $1.5 \cdot 10^{-6}$ | Rate of cell damage due to shear strain during cyclic loading | model fit |
| $k_{\mathrm{cl,fl}}$ $[\mathrm{s}^{-1}]$ | $1.5 \cdot 10^{-6}$ | Rate constant of aggrecan depletion due to fluid flow | model fit |
| $k_{\mathrm{aga}}$ [mol] | $0.250 \cdot 10^{-21}$ | Aggrecanase release from damaged cells | [8], model fit |
| $\alpha_{\mathrm{aga}}$ $[\mathrm{s}^{-1}]$ | $0.4 \cdot 10^{-5}$ | Rate constant to build up aggrecanase stimulus | [8] |
| $k_1$ $[\mathrm{s}^{-1}]$ | $3.5856 \cdot 10^{-5}$ | Rate constant for generating aggrecanases based on damaged cell-driven aggrecanase stimulus | [8] |

| | | | |
|---|---|---|---|
| $k_2$ $[\mathrm{s}^{-1}]$ | $10^{-4}$ | Aggrecanase degradation rate | [8,20] |
| $k_3$ $[\mathrm{s}^{-1}]$ | 0.9 | Catalytic rate constant for aggrecanase to degrade aggrecan | [8] |
| $K_{\mathrm{m,aga}}$ $[\mathrm{mol} \cdot \mathrm{m}^{-3}]$ | $5.5 \cdot 10^{-5}$ | Michaelis constant for aggrecanase | [8,21,22] |
| $P_{\mathrm{ag}}$ $[\mathrm{mol} \cdot \mathrm{cell}^{-1} \cdot \mathrm{s}^{-1}]$ | $2.4 \cdot 10^{-22}$ | Basal aggrecan biosynthesis rate (from healthy cells) | [8,23,24] |
| $C_{\mathrm{agg,tar}}$ $[\mathrm{mol} \cdot \mathrm{m}^{-3}]$ | 0.011635<br>Obtained from Kar *et al.*[8] as<br>$C_{\mathrm{agg,tar}} = \frac{C_{\mathrm{agg,tar,Kar}}}{\max(C_{\mathrm{agg,init,Kar}})} \cdot \max\,(C_{\mathrm{agg,init}}) = \frac{0.024}{0.0243} \cdot 0.01178$ | Target homeostatic aggrecan concentration | [8], recalibrated to [2] to fit free-swelling control model to data |
| $h_{\mathrm{z,agg}}$ $[\mathrm{m} \cdot \mathrm{s}^{-1}]$ | $2.7034 \cdot 10^{-10}$<br>Obtained from Kar *et al.*[8] as<br>$h_{\mathrm{z,agg}} = h_{\mathrm{z,agg,Kar}} \frac{\min(C_{\mathrm{agg,init,Kar}})}{\min(C_{\mathrm{agg,init}})} = 0.8 \cdot 10^{-10} \cdot \frac{0.0098}{0.0029}$ | Aggrecan axial mass transfer coefficient | [8], recalibrated to [2] |
| $h_{\mathrm{r,agg}}$ $[\mathrm{m} \cdot \mathrm{s}^{-1}]$ | $2.9034 \cdot 10^{-10}$<br>$h_{\mathrm{r,agg}} = h_{\mathrm{z,agg}} + 0.2 \cdot 10^{-10}$<br>addition as in Kar *et al.*[8] | Aggrecan lateral mass transfer coefficient | [8], recalibrated to [2] |
| $C_{\mathrm{agg,b}}$ $[\mathrm{mol} \cdot \mathrm{m}^{-3}]$ | 0 | Culture medium aggrecan concentration | [8] |

### References


1. Wilson, W., van Donkelaar, C., van Rietbergen, B. & Huiskes, R. A fibril-reinforced poroviscoelastic swelling model for articular cartilage. *J Biomech* **38**, 1195–1204 (2005).

2. Orozco, G., Tanska, P., Florea, C., Grodzinsky, A. & Korhonen, R. A novel mechanobiological model can predict how physiologically relevant dynamic loading causes proteoglycan loss in mechanically injured articular cartilage. *Sci Rep* **8**, 15599 (2018).

3. Mow, V. C. & Guo, X. E. Mechano-Electrochemical Properties Of Articular Cartilage: Their Inhomogeneities and Anisotropies. *Annu Rev Biomed Eng* **4**, 175–209 (2002).

4. Eskelinen, A. S. A. *et al.* Mechanobiological model for simulation of injured cartilage degradation via proinflammatory cytokines and mechanical stimulus. *PLoS Comput Biol* **16**, 1–25 (2020).

5. Saarakkala, S. & Julkunen, P. Specificity of fourier transform infrared (FTIR) microspectroscopy to estimate depth-wise proteoglycan content in normal and osteoarthritic human articular cartilage. *Cartilage* **1**, 262–269 (2010).

6. Kosonen, J. P. *et al.* Injury-related cell death and proteoglycan loss in articular cartilage: Numerical model combining necrosis, reactive oxygen species, and inflammatory cytokines. *PLoS Comput Biol* **19**, e1010337 (2023).

7. Li, L. P., Buschmann, M. D. & Shirazi-Adl, A. A fibril reinforced nonhomogeneous poroelastic model for articular cartilage: Inhomogeneous response in unconfined compression. *J Biomech* **33**, 1533–1541 (2000).

8. Kar, S. *et al.* Modeling IL-1 induced degradation of articular cartilage. *Arch Biochem Biophys* **594**, 37–53 (2016).

9. Quinn, T. M., Maung, A. A., Grodzinsky, A. J., Hunziker, E. B. & Sandy, J. D. Physical and biological regulation of proteoglycan turnover around chondrocytes in cartilage explants. Implications for tissue degradation and repair. *Ann N Y Acad Sci* **878**, 420–441 (1999).

10. Riegger, J. & Brenner, R. E. Pathomechanisms of posttraumatic osteoarthritis: Chondrocyte behavior and fate in a precarious environment. *Int J Mol Sci* **21**, (2020).

11. Momin, A., Perrotti, S. & Waldman, S. D. The role of mitochondrial reactive oxygen species in chondrocyte mechanotransduction. *Journal of Orthopaedic Research* (2023) doi:10.1002/jor.25709.

12. Eskelinen, ASA. *et al.* Cyclic loading regime considered beneficial does not protect injured and interleukin-1-inflamed cartilage from post-traumatic osteoarthritis. *J Biomech* **141**, 111181 (2022).

13. Orozco, G. A. *et al.* Shear strain and inflammation-induced fixed charge density loss in the knee joint cartilage following ACL injury and reconstruction: a computational study. *Journal of Orthopaedic Research* **40**, 1505–1522 (2022).

14. Argote, P. F. *et al.* Chondrocyte viability is lost during high-rate impact loading by transfer of amplified strain, but not stress, to pericellular and cellular regions. *Osteoarthritis Cartilage* **27**, 1822–1830 (2019).

15. Orozco, G. A. *et al.* Prediction of local fixed charge density loss in cartilage following ACL injury and reconstruction: A computational proof-of-concept study with MRI follow-up. *Journal of Orthopaedic Research* **39**, 1–8 (2020).

16. Hall, A. C., Urban, J. P. G. & Gehl, K. A. The effects of hydrostatic pressure on matrix synthesis in articular cartilage. *Journal of Orthopaedic Research* **9**, 1–10 (1991).

17. Elder, B. D. & Athanasiou, K. A. Hydrostatic Pressure in Articular Cartilage Tissue Engineering: From Chondrocytes to Tissue Regeneration. *Tissue Engineering: Part B* **15**, 1–12 (2009).

18. Morrell, K. C., Andrew Hodge, W., Krebs, D. E. & Mann, R. W. *Corroboration of in Vivo Cartilage Pressures with Implications for Synovial Joint Tribology and Osteoarthritis Causation*. www.pnas.orgcgidoi10.1073pnas.0507117102 (2005).

19. Loening, A. M. *et al.* Injurious mechanical compression of bovine articular cartilage induces chondrocyte apoptosis. *Arch Biochem Biophys* **381**, 205–212 (2000).

20. Yamamoto, K. *et al.* Low density lipoprotein receptor-related protein 1 (LRP1)-mediated endocytic clearance of a disintegrin and metalloproteinase with thrombospondin motifs-4 (ADAMTS-4): Functional differences of non-catalytic domains of ADAMTS-4 and ADAMTS-5 in LRP1 binding. *Journal of Biological Chemistry* **289**, 6462–6474 (2014).

21. Hooper, N. M. & Lendeckel, U. The ADAM family of proteases. in *Proteases in Biology and Disease* 1–30 (2005). doi:10.1007/b106833.

22. Wittwer, A. J. *et al.* Substrate-dependent inhibition kinetics of an active site-directed inhibitor of ADAMTS-4 (aggrecanase 1). *Biochemistry* **46**, 6393–6401 (2007).

23. Zhang, L., Gardiner, B. S., Smith, D. W., Pivonka, P. & Grodzinsky, A. A fully coupled poroelastic reactive-transport model of cartilage. *MCB Molecular and Cellular Biomechanics* **5**, 133–153 (2008).

24. Sengers, B. G., Taylor, M., Please, C. P. & Oreffo, R. O. C. Computational modelling of cell spreading and tissue regeneration in porous scaffolds. *Biomaterials* **28**, 1926–1940 (2007).

190
